# Growth Factor-Independent mTORC1 Signaling Promotes Primary Cilia Length via Suppression of Autophagy

**DOI:** 10.1101/2025.05.07.652626

**Authors:** Jong Shin, Yann Cormerais, Khaled Tighanimine, Pratima Niroula, Madi Y. Cisse, Samuel C. Lapp, Krystle C. Kalafut, Sandra Schrötter, Brendan D. Manning

## Abstract

The mechanistic target of rapamycin (mTOR) complex 1 (mTORC1), as a sensor of growth signals that subsequently controls cell growth, has been predominantly studied in actively proliferating cells. Primary cilia are sensory organelles present on most quiescent cells, where they play essential roles in receiving environmental and developmental signals. Given that ciliated cells are non-proliferative, we investigated whether mTORC1 signaling influences the growth of primary cilia. Here, we show that mTORC1 promotes primary cilia elongation, without effects on ciliogenesis or cell growth, by suppressing autophagy. Inhibition of mTORC1 signaling through pharmacological, nutritional, or genetic interventions gave rise to shortened primary cilia, while activation of the pathway resulted in elongation. Furthermore, pharmacological or genetic inhibition of autophagy, a key downstream process blocked by mTORC1, also elongated primary cilia and rendered them resistant to mTORC1 inhibition. Notably, these mTORC1-mediated effects on primary cilia extend to mouse neurons ex vivo and in vivo. These findings highlight a previously unrecognized role for mTORC1 signaling in the control of primary cilia length that may contribute to diseases where ciliary function is altered, referred to as ciliopathies.

## INTRODUCTION

The mechanistic target of rapamycin (mTOR) complex 1 (mTORC1) is a central regulator of cellular metabolism and growth, integrating upstream nutrient and growth factor signals to drive anabolism while suppressing catabolic processes^1,2^. The vast majority of studies on mTORC1 function have been in proliferating cells stimulated with growth factors, leaving questions as to the specific role of mTORC1 regulation by nutrients, which are essential for the basal growth factor-independent activity of mTORC1.

While it is likely that mTORC1 senses many essential nutrients, the mechanisms by which it senses changes in specific amino acids are best understood^3^. Amino acids activate mTORC1 through the Rag GTPases, which function as heterodimers of RagA or B bound to RagC or D^4^. In the presence of amino acids, the heterodimer is in the RagA/B^GTP^-RagC/D^GDP^ conformation that can directly bind to mTORC1 and recruit it to the outer lysosomal membrane, where it can be further activated^5^. Amino acid availability influences the activation state of a protein complex comprised of NPRL2, NPRL3, and DEPDC5, referred to as GATOR1, which in the absence of amino acids functions as a GTPase-activating protein (GAP) for RagA/B, thus promoting its conversion to the GDP-bound state unable to engage mTORC1^6,7^. The amino acid-dependent recruitment of mTORC1 to the lysosome through the Rag GTPases is essential for both the basal activation of mTORC1 and its ability to be regulated by growth factors^2^. In its GTP-bound state, the RHEB GTPase binds to mTORC1 recruited to the lysosomal surface and stimulates its kinase activity^8^. A key negative regulator of RHEB and its activation of mTORC1 is the tuberous sclerosis complex (TSC) protein complex, comprised of TBC1D7, TSC1, and TSC2, which acts as a GAP for Rheb^9–15^. The TSC complex receives inhibitory signals from growth factor signaling pathways, and loss of its constituent components leads to elevated Rheb-GTP loading and growth factor-independent activation of mTORC1^9,16–21^.

In addition to the tumor and neurological syndrome TSC, dysregulation of mTORC1 signaling is implicated in numerous pathological conditions, including cancer, metabolic disorders such as diabetes and obesity, and neurological disorders such as epilepsy^1,22^. Mechanistically, uncontrolled mTORC1 signaling drives aberrant cellular proliferation, metabolic changes, and impaired physiological responses, contributing to disease onset and progression. Conversely, insufficient mTORC1 signaling in specific settings can disrupt cellular growth, weaken immune responses, and impair differentiation. Given the wide-ranging effects of its dysregulation in pathological settings, understanding the full suite of downstream functions for mTORC1 signaling is crucial.

The activation state of mTORC1 influences the metabolic balance of cells between anabolic and catabolic states^1,2^. mTORC1 stimulates key anabolic processes, including the *de novo* synthesis of proteins, lipids, and nucleotides, in part, through its canonical downstream effectors the ribosomal S6 kinases (S6K1 and S6K2) and the eIF4E-binding proteins (4E-BP1 and 4E-BP2), thereby supporting the production of biomass to drive cell growth. Reciprocally, mTORC1 suppresses catabolic processes, including bulk autophagy—a conserved intracellular degradation system that recycles organelles and other cytosolic contents to restore intracellular nutrients to nutrient deprived cells^23^. Thus, autophagy is acutely induced upon inhibition of mTORC1 signaling in response to either pharmacological inhibitors or nutrient depletion.

Much of the research on mTORC1 signaling has focused on proliferating cells, extensively characterizing its role in promoting cell growth and division—processes that demand high energy and nutrient resources for macromolecular synthesis and organelle duplication. However, its function in non-proliferative cells remains less understood. The vast majority of cells in the body exist in a quiescent or terminally differentiated non-proliferative state, yet in many tissues mTORC1 signaling remains active and can continue to fluctuate in response to diurnal cycles, developmental cues, nutrient availability, and immune challenges^1^. To function effectively in these conditions, non-proliferative cells must allocate resources efficiently for organellar, cellular, and organismal homeostasis. Unlike proliferating cells, which engage mTORC1 signaling to drive anabolism in support of biomass production for growth and division, the role of mTORC1 in non-proliferative cells is more poorly understood.

Primary cilia are highly organized microtubule-based sensory organelles predominantly present on non-proliferative, terminally differentiated cells^24,25^. Depending on the cellular context, primary cilia propagate developmental or mechanical signals to cells^26^. The critical role of these organelles extending from the cell surface is highlighted by ciliopathies—a group of severe genetic disorders caused by mutations in ciliary components, leading to a wide range of developmental and metabolic abnormalities^26^. Notably, ciliogenesis is tightly linked to the cell cycle, with cilia typically assembling when cells exit the cell cycle and enter a quiescent state^27–30^. In non-transformed cells, growth factor withdrawal leads to cell cycle arrest and quiescence, a cellular state that promotes ciliogenesis^31,32^.

In many differentiated and quiescent cells, primary cilia co-exist with basal mTORC1 signaling, raising the question of whether mTORC1 activity influences ciliary architecture. While a few previous studies have examined this relationship, the conclusions have been inconsistent. It has been found that mTORC1 activation promotes the lengthening of primary cilia in zebrafish embryos^33^. In contrast, mTORC1 inhibition with rapamycin was found to increase the length of primary cilia in cultured mammalian renal epithelial and vascular endothelial cells^34^. Additional complexity arises from studies involving loss of TSC1 or TSC2, which leads to growth factor-independent mTORC1 activation. One study found that MEFs lacking either *Tsc1* or *Tsc2* are more likely to be ciliated and that their cilia are longer, with rapamycin having no effect on this increased length^35^. A second study observed ciliary elongation in *Tsc1*-null MEFs that was reversible with rapamycin, whereas, paradoxically, an independent line of *Tsc2*-null MEFs exhibited shortened primary cilia and reduced overall ciliogenesis^36^. Finally, it has been shown there is a reduction in the number of ciliated neurons in cortical tubers, the most common brain lesion in TSC patients^37^. Furthermore, a mouse model of neuronal *Tsc2* loss exhibited a reduction in ciliated neurons, which could be rescued by rapamycin, although rapamycin did not enhance ciliogenesis in wild-type neurons^37^. These discrepancies might reflect the use of non-isogenic cells, variations in cell types, or differences in the nature and degree of mTORC1 perturbation.

In this study, we investigate the potential role of mTORC1 signaling in controlling primary cilia dynamics. Using complementary pharmacological and genetic approaches across isogenic models, we determined the effects of mTORC1 activation and inhibition on primary cilia formation and length under nutrient-sensitive but growth factor–independent conditions. Our findings reveal that mTORC1 promotes primary cilia elongation through the suppression of autophagy, independent of effects on cell proliferation or growth.

## RESULTS

### mTORC1 inhibition shortens primary cilia

Primary cilia formation is closely tied to cell cycle progression^27–30^. We first aimed to establish a reliable kinetic profile of primary cilia formation using the human retinal pigment epithelial cell line (RPE1), an hTERT-immortalized, non-transformed human cell line widely studied for primary cilia biology. Serum withdrawal from the culture medium induced the accumulation of cells in the G0/G1 phase of the cell cycle by 24 hours, with ciliated cells appearing by 12 hours (Figure S1A-C)^38,39^. We observed a progressive increase in the number of ciliated cells and a corresponding elongation of primary cilia to a maximum by 24 hours of serum withdrawal. The appearance of ciliated cells coincided with a progressive accumulation of the cell cycle-exited G0 population, as indicated by accumulation of a p27 mutant (p27K-) that lacks CDK binding (Figure S1D)^40^. By 24 hours of serum withdrawal, the majority of cells had entered G0, and the number of ciliated cells and primary cilia length remained relatively unchanged thereafter, suggesting that an equilibrium between primary cilia assembly and disassembly had been reached (Figure S1B-D). This 24-hour time point was thus selected for subsequent assessments of the effect of mTORC1 signaling on primary cilia dynamics.

Serum readdition to cells that became quiescent after 24 hours of serum starvation triggered cell cycle re-entry, an increase in total cell number, and an increase in cell size (Figure 1A-C). Consistent with growth factor-independent mTORC1 signaling not influencing the quiescent state of RPE1 cells in the absence of serum, treatment with the mTOR inhibitors rapamycin or Torin1, a potent ATP-competitive inhibitor of mTOR, resulted in only minimal changes in cell number, cell-cycle distribution, and cell size (Figure 1A-C)^41,42^. These data indicate that the canonical anabolic effects of mTORC1 signaling that promote cell proliferation and cell growth are predominantly exerted in actively proliferating, growth factor-stimulated cells and dispensable in quiescent cells.

**Figure 1.**
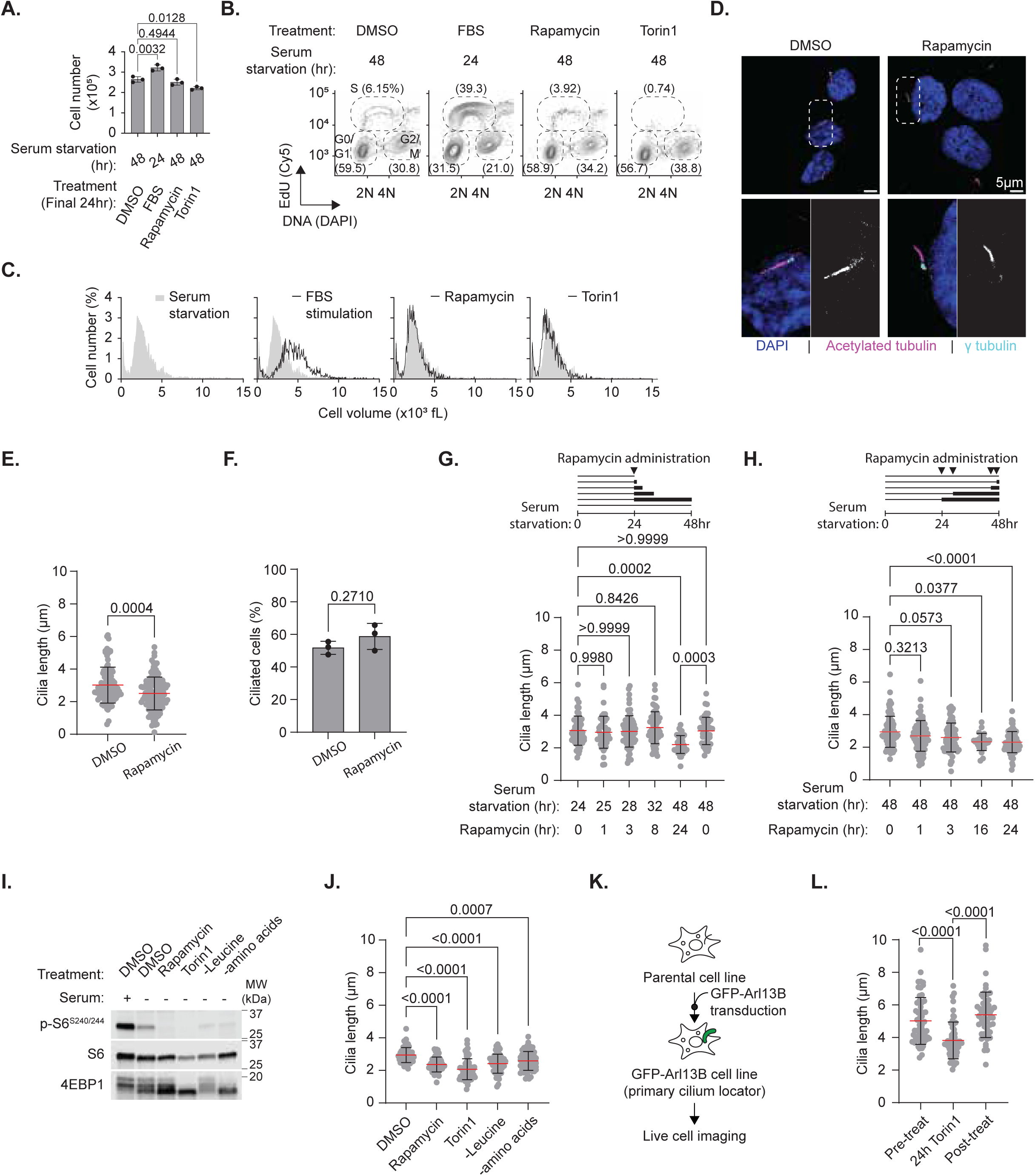
mTORC1 inhibition shortens primary cilia. (A–C) (A) Cell number, (B) flow cytometric analysis of cell cycle distribution via genomic incorporation of EdU as a function of DAPI-based nuclear intercalation, and (C) distribution of cell volumes in RPE1 cells treated with DMSO, FBS (10%), rapamycin, or Torin1 for the indicated times. (D) Confocal microscopy images of the immunofluorescent axoneme (acetylated tubulin in magenta) and basal body (γ-tubulin in cyan) of primary cilia in RPE1 cells treated with DMSO (left) or rapamycin (right) for the final 24 hours. Regions of interest (dotted in top panels) are shown at higher magnification in the bottom panels as merged images (left) and single-channel grayscale for acetylated tubulin (right). Nuclei are stained with DAPI. Scale bar: 5 μm. (E–F) (E) Primary cilia length and (F) percentage of ciliated cells in RPE1 cells treated with DMSO or rapamycin. (G) Time-course analysis of primary cilia length in RPE1 cells treated with rapamycin after 24-hour serum starvation. (H) Primary cilia length in RPE1 cells treated with rapamycin for varying durations within a total of 48 hours of serum starvation. (I–J) (I) Western blotting and (J) primary cilia length of RPE1 cells treated with DMSO, rapamycin, Torin1, leucine deprivation, or complete amino acid deprivation for the final 24 hours in serum-free conditions (48 h total). (K) Schematic diagram of the generation of the GFP-Arl13B RPE1 cell line and live-cell imaging for primary cilia length measurement. (L) Primary cilia length of GFP-Arl13B RPE1 cells before Torin1 treatment, after 24 h of treatment with Torin1 (250 nM), and 24 h after Torin1 removal. All treatments were applied for the final 24 hours in serum-free conditions (48 h total), unless otherwise specified: DMSO (0.1%), rapamycin (20 nM), Torin1 (250 nM). Statistical analysis was performed using two-way ANOVA followed by Šídák’s multiple comparisons test. The significance threshold (α) was set at 0.05, and all p-values are reported.

Treatment of ciliated cells with rapamycin for 24 hours resulted in a significant reduction in primary cilia length without affecting the number of ciliated cells in the population (Figure 1D-F). Importantly, although mTORC1 signaling, as measured by phosphorylation of S6K1, its substrate ribosomal S6, and 4EBP1 (by mobility shift)—canonical downstream effectors of mTORC1—is reduced in the absence of serum compared to serum-fed conditions, it remains sensitive to rapamycin under these conditions that promote cell ciliation (Figure S1E). To assess the kinetics of rapamycin-induced primary cilia shortening, we measured primary cilia length under two experimental conditions: (1) time-dependent analysis of the drug effect following 24 hours of serum starvation (Figure 1G), and (2) varying durations of rapamycin treatment within a total of 48 hours of serum starvation (Figure 1H). Both approaches demonstrated that rapamycin induces primary cilia shortening in a time-dependent manner. Furthermore, we confirmed that rapamycin significantly reduced primary cilia length across three widely used culture media formulations optimized for mammalian cell culture (Figure S1F).

We next explored additional pharmacological and nutritional strategies to inhibit mTORC1 signaling. Suppression of mTORC1 by treatment with Torin1 or by withdrawal of either all amino acids or leucine alone led to significantly shorter primary cilia, with the degree of shortening largely correlating the decrease in markers of mTORC1 signaling (Figure 1I, J). This measured reduction in primary cilia length did not correlate with changes in the G0 population (Figure S1G). Additionally, consistent with the finding that mTOR inhibitors did not decrease cell size under culture conditions promoting ciliogenesis (Figure 1C), neither rapamycin nor Torin1 decreased total cellular protein content under these conditions, as measured with carboxyfluorescein succinimidyl ester (CFSE) (Figure S1H)^43^.

To further validate these findings using an alternative approach to visualize primary cilia in live cells, we generated a stable RPE1 cell line expressing GFP-conjugated Arl13B, a small GTPase highly enriched in the membrane of primary cilia (Figure 1K)^44,45^. We measured GFP-labelled primary cilia in live cells at three time points: before Torin1 treatment, after 24 hours of Torin1 treatment, and 24 hours after Torin1 removal. Consistent with our immunofluorescence findings, we observed that Torin1 treatment significantly shortened primary cilia, while removal of the drug led to elongation of cilia back to their original lengths, thereby demonstrating the reversibility of this effect (Figure 1L). Collectively, these data demonstrate that inhibition of mTORC1 signaling leads to primary cilia shortening, suggesting that changes in nutrient levels, such as amino acids, can modulate primary cilia length through the regulation of mTORC1 signaling.

### Genetic manipulation of mTORC1 signaling demonstrates that it promotes the elongation of primary cilia

While the shortening of primary cilia upon pharmacological or nutrient-mediated inhibition of mTORC1 signaling suggested that the pathway controls primary cilia length, we sought to confirm its role via genetic perturbations of the pathway. We generated stable RPE1 cell lines with doxycycline-inducible expression of a short-hairpin RNA specifically targeting the *MTOR* gene or non-targetting scrambled control shRNA. A substantial reduction in mTOR protein expression and phosphorylation of the direct mTORC1 target S6K1 was observed specifically upon treatment of the shMTOR cells with doxycycline (Figure 2A). This attenuation of mTOR signaling coincided with a significant decrease in primary cilia length (Figure 2B), recapitulating effects observed with pharmacological or nutrient-mediated inhibition of mTORC1.

**Figure 2.**
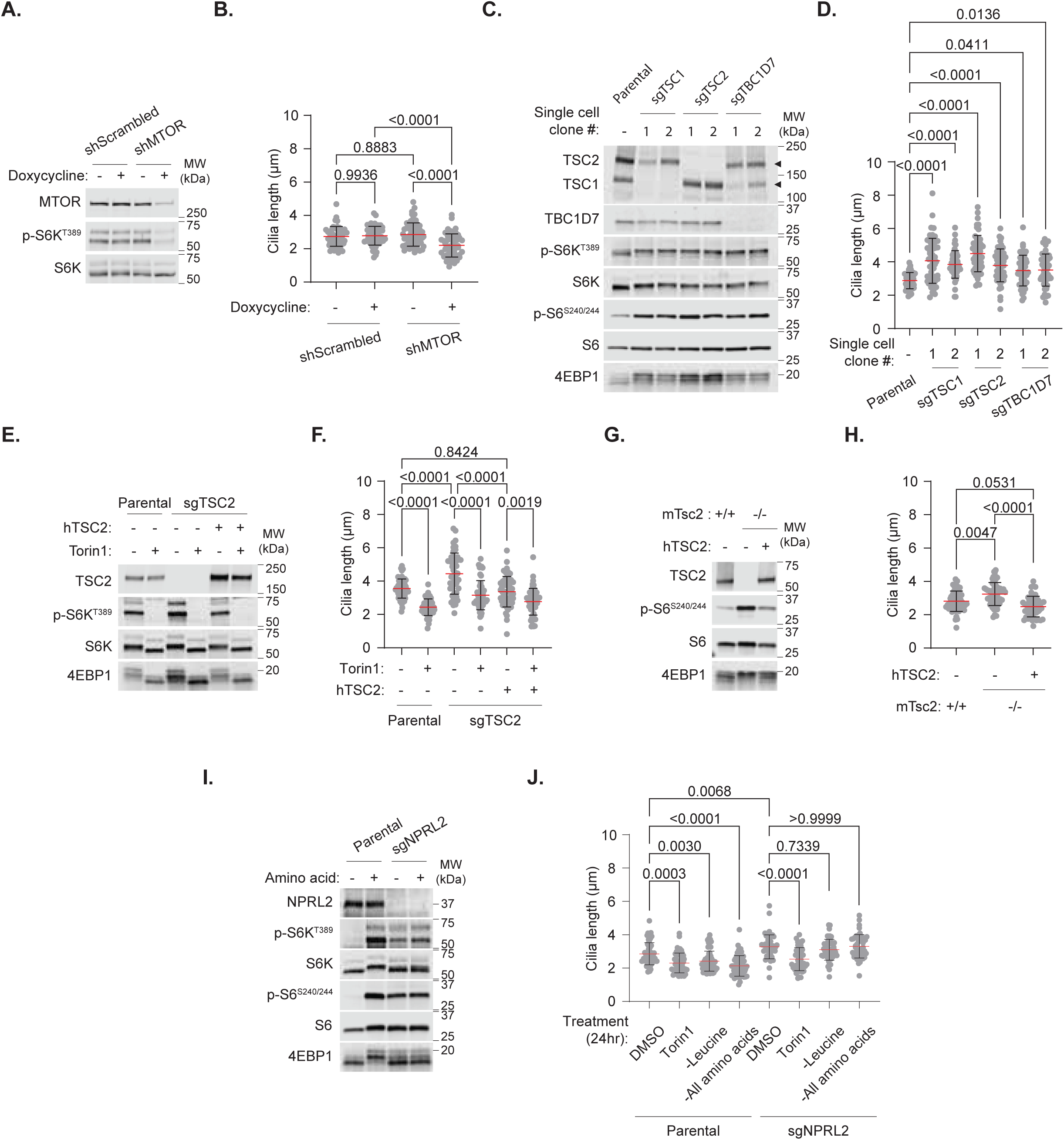
Genetic manipulation of mTORC1 signaling demonstrates that it promotes the elongation of primary cilia. (A–B) (A) Immunoblot and (B) primary cilia length in doxycycline-inducible shMTOR RPE1 cells treated with doxycycline (0.5 µg/mL) for 7 days. (C) Immunoblot of clonal RPE1 cell lines with sgRNAs targeting TSC1, TSC2, or TBC1D7. Cells were serum-starved overnight. (D) Primary cilia length in RPE1 lines used in (C). TSC1 and TSC2 are indicated by black arrowheads. (E) Immunoblot of sgTSC2 RPE1 cells, human TSC2-rescued sgTSC2 cells, and parental controls treated with Torin1 for 1 hour in growth media with serum. (F) Primary cilia length in RPE1 cell lines used in (E) following Torin1 treatment. (G) Immunoblot of *Tsc2^+/+^*, *Tsc2^−/−^*, and human TSC2-rescued *Tsc2^−/−^* MEFs after overnight serum starvation. (H) Primary cilia length in MEF cell lines used in (G). (I) Immunoblot of parental and sgNPRL2 RPE1 cells treated with overnight amino acid starvation followed by 1 hour of amino acid stimulation. (J) Primary cilia length in parental and sgNPRL2 RPE1 cells treated with DMSO, Torin1, leucine deprivation, or complete amino acid deprivation. All treatments for primary cilia assessment were applied during the final 24 hours in serum-free conditions (48 h total): DMSO (0.1%), rapamycin (20 nM), Torin1 (250 nM). Statistical analysis was performed using two-way ANOVA followed by Šídák’s multiple comparisons test. The significance threshold (α) was set at 0.05, and all p-values are reported.

We next sought to determine whether an elevation in mTORC1 signaling through deletion of TSC complex components would lead to primary cilia elongation. However, deletion of *TSC1* or *TSC2* in non-transformed cells is known to induce permanent cell cycle arrest (or senescence) that is dependent on p53 (*TP53*), limiting our ability to assess effects on primary cilia^17^. To bypass this, the CRISPR-Cas9 gene editing system was used to delete a TSC complex component (sgTSC1, sgTSC2, sgTBC1D7) in RPE1 cells where p53 had already been deleted (sgTP53)^17^. After confirming p53 deletion, as evidenced by the absence of p53 protein accumulation upon Actinomycin D treatment—a canonical DNA damage response—we subsequently deleted TSC complex components and confirmed that these cells bypassed p53-mediated cell-cycle arrest (Figure S2A, B). While p53 deletion alone did not significantly alter mTORC1 signaling (Figure S2C), additional deletion of a TSC complex component in sgTP53 cells constitutively elevated mTORC1 signaling, even in the absence of serum (Figure 2C). Importantly, while the formation and elongation of primary cilia in sgTP53 cells were comparable to those in parental cells (Figure S2D, E), all isogenic TSC-null cell lines tested exhibited significantly longer primary cilia (Figure 2D). Inhibition of mTORC1 signaling with Torin1 shortened primary cilia in both parental and sgTSC2 cells, as did stable re-expression of a human TSC2 cDNA (Figure 2E, F). To confirm the effects of TSC2 loss on primary cilia length in a distinct cell type and species, we employed an established *Tsc2*-null mouse embryo fibroblast (MEF) cell line (p53-null)^17^. Indeed, *Tsc2*-null MEFs displayed longer primary cilia than their littermate-derived wild-type counterparts, an effect rescued by stable re-expression of human TSC2 (Figure 2G, H).

We next tested whether the primary cilia-shortening effects of amino acid withdrawal were dependent on mTORC1 inhibition. To test this, we genetically deleted *NPRL2* (sgNPRL2), a key component of GATOR1, leading to mTORC1 activation even in the absence of amino acids (Figure 2I)^46^. Unlike parental cells, primary cilia lengths remained resistant to leucine or amino acid deprivation in sgNPRL2 cells but were still shortened by Torin1 (Figure 2J). Collectively, these data establish that mTORC1 signaling promotes primary cilia elongation.

### mTORC1 signaling promotes elongation of primary cilia via autophagy suppression

As autophagy is one of the only processes downstream of mTORC1 that is robustly induced by nutrient deprivation but more resistant to growth factor withdrawal, we sought to determine whether its regulation might contribute to the mTORC1-mediated effects on primary cilia^2^. To test this, an RPE1 cell line expressing a GFP-LC3B-RFP fusion protein as a reporter of autophagic flux was generated, enabling fluorescent visualization of GFP-conjugated LC3B degradation in lysosomes as a readout of autophagic activity, while RFP, cleaved from the GFP-LC3B-RFP by ATG4 activation, serves as an internal control (Figure 3A)^47^. In this system, a high GFP/RFP ratio indicates reduced autophagic flux, whereas a low ratio signifies active autophagic flux. Using this reporter cell line, we first confirmed that mTORC1 inhibition by rapamycin, Torin1, or leucine deprivation increased autophagy flux, as shown by a significant decrease in the GFP/RFP ratio (Figure 3B, C). Conversely, treatment with the vacuolar ATPase inhibitor bafilomycin A1, which inhibits lysosomal acidification and functions, including autophagy, led to an increased GFP/RFP ratio (Figure 3B, C). These results confirm that mTORC1 signaling negatively regulates autophagy flux in this setting.

**Figure 3.**
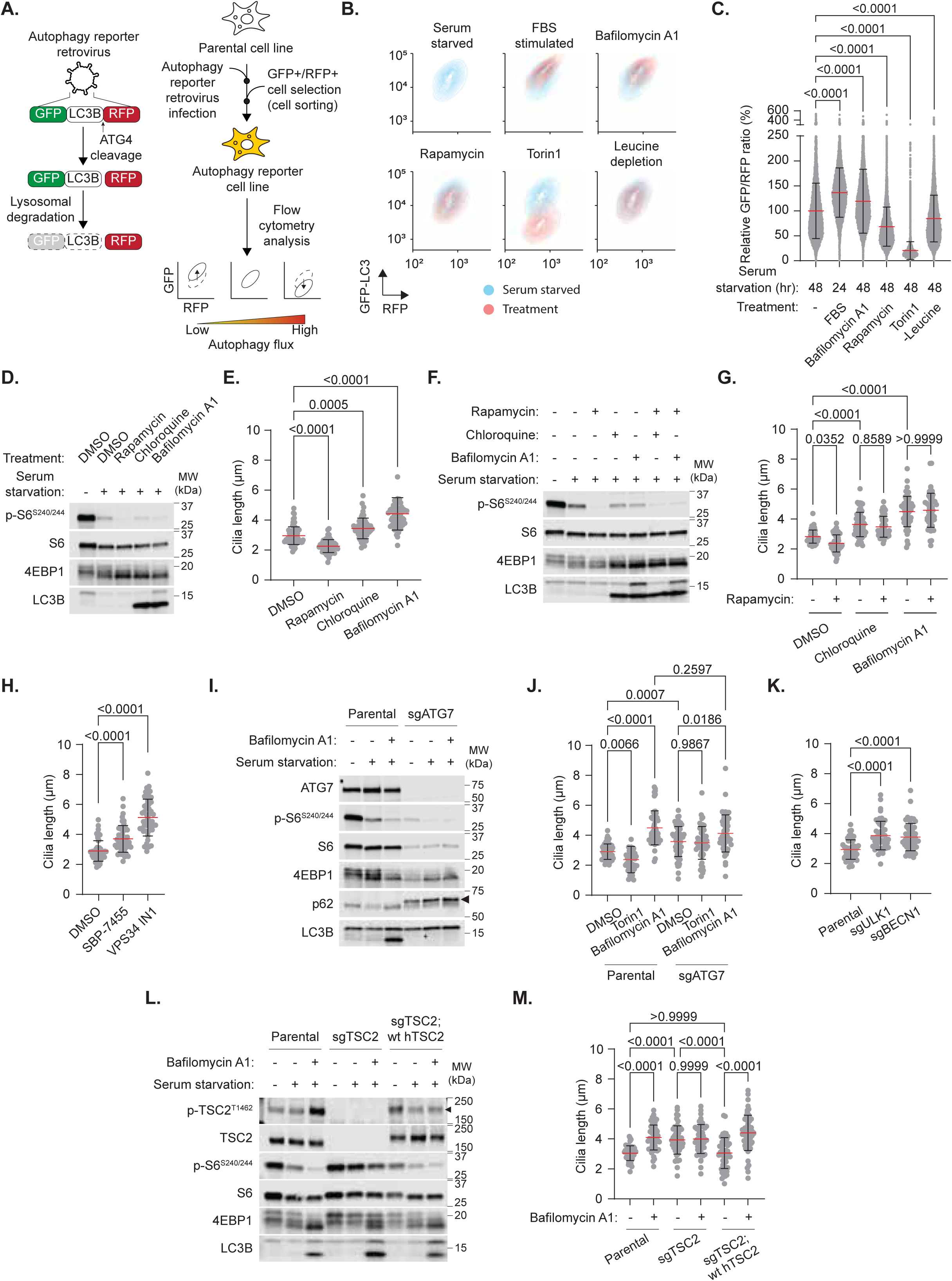
mTORC1 signaling promotes elongation of primary cilia via autophagy suppression. (A) Schematic diagram showing the generation of the GFP-LC3B-RFP autophagy reporter RPE1 cell line and flow cytometry analysis used to assess autophagic flux. (B–C) (B) Flow cytometric analysis and (C) quantification of autophagic flux in RPE1 cells treated with FBS (10%), bafilomycin A1, rapamycin, Torin1, or leucine deprivation for the indicated times. (D) Immunoblot and (E) primary cilia length in RPE1 cells treated with DMSO, rapamycin, chloroquine, or bafilomycin A1. (F–G) (F) Immunoblot and (G) primary cilia length in RPE1 cells treated with DMSO, chloroquine, or bafilomycin A1, alone or in combination with rapamycin. (H) Primary cilia length in RPE1 cells treated with DMSO, ULK1/2 inhibitor (SBP-7455), or VPS34 inhibitor (VPS34-IN1). (I) Immunoblot of parental and sgATG7 RPE1 cells treated with bafilomycin A1 for the final 3 hours under overnight serum starvation. (J) Primary cilia length in RPE1 cells used in (I) following treatment with DMSO, Torin1, or bafilomycin A1. p62 is indicated by a black arrowhead. (K) Primary cilia length in parental, sgULK1, or sgBECN1 RPE1 cells. (L) Immunoblot of parental, sgTSC2, and human TSC2-rescued sgTSC2 RPE1 cells treated with bafilomycin A1 for the final 3 hours under overnight serum starvation. (M) Primary cilia length in RPE1 cells used in (L) following treatment with DMSO or bafilomycin A1. All treatments were applied during the final 24 hours in serum-free conditions (48 h total), unless otherwise specified: DMSO (0.1%), rapamycin (20 nM), Torin1 (250 nM), chloroquine (50 µM), bafilomycin A1 (100 nM), SBP-7455 (10 µM), VPS34-IN1 (10 µM). Statistical analysis was performed using two-way ANOVA followed by Šídák’s multiple comparisons test. The significance threshold (α) was set at 0.05, and all p-values are reported.

To determine the effects of autophagy on primary cilia, we tested whether autophagy inhibition affects primary cilia length. Treatment with bafilomycin A1 or chloroquine, chemically distinct inhibitors of the lysosome, blocked the autophagic degradation of lipidated LC3B (lower band in Figure 3D). In contrast to rapamycin, treatment with these compounds led to a significant elongation of primary cilia (Figure 3E). Notably, co-treatment with rapamycin and either lysosomal inhibitor prevented the rapamycin-induced shortening of primary cilia (Figure 3F, G), suggesting that mTORC1 inhibition shortens primary cilia in a lysosome-dependent manner. Since bafilomycin A1 and chloroquine are known to inhibit the later stages of autophagy, we sought to block the earlier steps of autophagy to further determine its role in controlling primary cilia length^48,49^. Mechanistically, mTORC1 has been established to suppress the initiation of autophagy through the inhibition of ULK1^50–52^. To this end, we observed that pharmacological inhibition of ULK1/2 with the selective inhibitor SBP-7455 led to primary cilia elongation (Figure 3H)^53^. In addition, we observed a pronounced effect with VPS34 IN1, a selective inhibitor of VPS34, the class III PI3K that facilitates autophagosome formation, further suggesting that autophagy negatively influences primary cilia length (Figure 3H)^54,55^.

We next sought to genetically disrupt genes essential for autophagy to further determine its role in the control of primary cilia length by mTORC1 signaling. *ATG7* was genetically deleted using CRISPR-Cas9 (sgATG7), leading to the absence of lipidated LC3B and accumulation of p62 (SQSTM1), two consequences confirming defects in autophagic flux (Figure 3I)^56^. Consistent with our findings using pharmacological inhibitors of autophagy, sgATG7 cells exhibited elongated primary cilia that were no longer shortened by Torin1 treatment (Figure 3J). A similar increase in primary cilia length was measured in cells with genetic deletion of two other autophagy genes, *ULK1* or *BECN1* (Figure 3K, S3A).

Reciprocal to the use of mTOR inhibitors, we measured autophagic flux in RPE1 cells with elevated mTORC1 signaling due to disruption of TSC complex components. Indeed, sgTSC1, sgTSC2, or sgTBC1D7 cells all showed significantly reduced autophagy flux relative to parental cells, whereas Torin1 induced autophagy similarly in all cells (Figure S3B, C), demonstrating an mTOR-dependent decrease in autophagic flux in these cells. Bafilomycin A1 treatment blocked autophagic flux regardless of TSC2 status in these cells (Figure 1L). However, while bafilomycin A1 treatment increased primary cilia length in parental cells, it failed to further elongate primary cilia in sgTSC2 cells, an effect rescued by re-expression of TSC2 (Figure 3M). These data support the conclusion that mTORC1 signaling elongates primary cilia through the suppression of autophagy.

### mTORC1 regulates primary cilia length in the mouse brain

To extend our findings to an *in vivo* context, we focused on the brain, where most neurons are post-mitotic and known to possess prominent primary cilia^57,58^. We first cultured primary cortical neurons from embryonic day 16.5 (E16.5) C57BL/6J embryos and treated them with mTORC1 inhibitors or bafilomycin A1 (Figure 4A). Consistent with our findings in cultured RPE1 cells and MEFs, treatment of primary neurons with Torin1 resulted in a significant reduction in primary cilia length, with a modest decrease observed with rapamycin treatment, whereas treatment with bafilomycin A1 led to increased primary cilia length (Figure 4B-D). These results suggest that the effects of mTORC1 signaling and autophagy on primary cilia extend to mouse cortical neurons.

**Figure 4.**
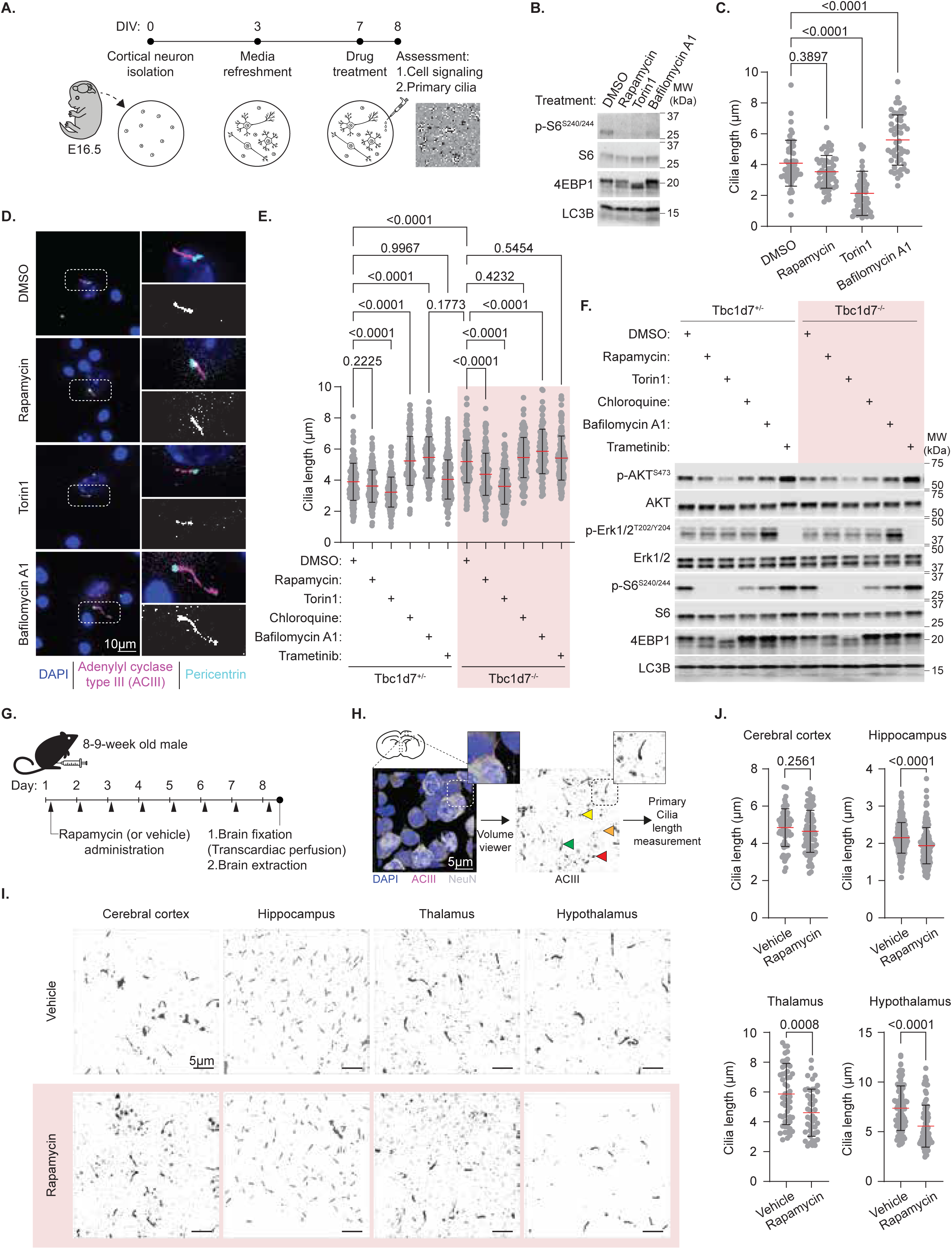
mTORC1 regulates primary cilia length in neurons. (A) Schematic overview of ex vivo experiments using cortical neurons isolated from E16.5 mouse embryos to assess primary cilia length changes in response to mTORC1 signaling and autophagy. (B–C) (B) Immunoblot and (C) primary cilia length in cortical neurons cultured from C57BL/6 E16.5 embryos and treated with DMSO, rapamycin, Torin1, or bafilomycin A1. (D) Confocal microscopy images of immunofluorescent axoneme (adenylyl cyclase type III, ACIII, in magenta) and basal body (pericentrin in cyan) of primary cilia in C57BL/6 cortical neurons treated with DMSO, rapamycin, Torin1, or bafilomycin A1 for 24 h. Dotted regions (left panel) are shown at higher magnification in the right panels, with merged image (top) and single channel grayscale for ACIII (bottom). Nuclei are stained with DAPI. (E–F) (E) Primary cilia length and (F) immunoblot in cultured *Tbc1d7^+/−^*and *Tbc1d7^−/−^* cortical neurons treated with DMSO, rapamycin, Torin1, chloroquine, bafilomycin A1, or trametinib. (G,H) Schematic overview of *in vivo* experiments using 8–9-week-old male C57BL/6 mice treated with daily intraperitoneal injections of rapamycin (1 mg/kg body weight) to assess primary cilia length changes in various brain regions. (H) Experimental overview showing volume viewer-mediated 3D rendering of Z-stack confocal immunofluorescent images of brain regions of interest. Primary cilia were visualized with adenylate cyclase type III, ACIII, (magenta), neurons with NeuN (grey), and nuclei with DAPI (blue). Dotted regions (top right corner) are shown at higher magnification in the right edge of each image. For primary cilia length assessment, only well-stained, tubular-shaped primary cilia (green arrows) were measured. Pixels excluded from analysis included those with high background (yellow), weak/discontinuous/unsuccessful staining (orange), or ambiguous morphology (red). (I-J) (I) 3D-rendered images and (J) quantification of primary cilia length from four distinct brain regions. All treatments of cultured neurons were applied for 24 h: DMSO (0.1%), rapamycin (20 nM), Torin1 (250 nM), chloroquine (50 µM), bafilomycin A1 (100 nM), trametinib (5 µM). Unpaired two-tailed Student’s t-test was used for comparisons between two groups. For comparisons between three or more groups, two-way ANOVA followed by Šídák’s multiple comparisons test was used. The significance threshold (α) was set at 0.05, and all p-values are reported.

To investigate whether a modest increase in mTORC1 signaling influences primary cilia length in primary neurons, cortical neurons were isolated from mouse embryos with the endogenous *Tbc1d7* gene heterozygously or homozygously deleted (Figure S4A)^59^. Unlike other TSC complex components, TBC1D7 is dispensable for embryonic development, but *Tbc1d7^−/−^* mice display enhanced brain mTORC1 signaling and postnatal brain growth^59^. To obtain genetically mixed littermate embryos, we crossed *Tbc1d7^+/−^*mice with *Tbc1d7^−/−^* mice (Figure S4B, C). Cortical neurons were cultured, as above, and treated with inhibitors of mTOR, the lysosome, or, as a negative control, the protein kinase MEK. *Tbc1d7* null primary cortical neurons possessed longer primary cilia than those of *Tbc1d7^+/−^* neurons from littermate embryos (Figure 4E, F). The elongated primary cilia of *Tbc1d7^−/−^* neurons were significantly shortened by treatment with rapamycin or Torin1, whereas bafilomycin A1 treatment increased primary cilia length only in *Tbc1d7^+/−^* neurons (Figure 4E). However, treatment with the MEK inhibitor trametinib did not alter primary cilia length in neurons of either genotype (Figure 4E, F).

Finally, to determine whether systemic treatment with mTORC1 inhibitors can influence the length of primary cilia in the mouse brain, we administered rapamycin (1 mg/kg) or vehicle daily, via intraperitoneal injection, for one week to adult (8–9-week-old) C57BL/6 male mice (Figure 4G). Sections from harvested brains were subjected to immunofluorescence imaging and 3D rendering of primary cilia using adenylyl cyclase type III (ACIII) staining (Figure 4H)^60^. Quantitative measurements of primary cilia length in four brain regions revealed a significant reduction in primary cilia length in the hippocampus, thalamus, and hypothalamus, but not the cerebral cortex, after rapamycin treatment (Figure 4I, J). Therefore, these findings confirm that mTORC1 signaling regulates primary cilia length in regions of the adult mammalian brain.

## DISCUSSION

This study reveals a role for mTORC1 signaling in the regulation of primary cilia length through its effects on autophagy. Inhibition of mTORC1 through pharmacological, nutrient, or genetic interventions shortened primary cilia, while increasing mTORC1 signaling resulted in primary cilia elongation. Furthermore, pharmacological or genetic inhibition of autophagy elongated primary cilia, and rendered them resistant to inhibition of mTORC1 signaling. These findings extend to mouse neurons ex vivo and to multiple regions of mouse brain in vivo.

The ability of mTORC1 signaling to influence primary cilia length highlights a link between nutrient sensing and the dynamic growth of this specialized signaling organelle. mTORC1 integrates numerous cellular signals and can be attenuated by various forms of cellular stress, including metabolic imbalance, energy deficiency, proteostatic disruption, and genomic instability ^61–63^. In turn, it coordinates the synthesis and recycling of macromolecules in support of cellular homeostasis. To the diverse set of known mTORC1 functions, the findings reported here add the control of primary cilia length, merging its regulation to the many changes in cellular state sensed by mTORC1. In quiescent or differentiated cells, this regulation may help calibrate the sensory and signaling capacity of primary cilia to meet physiological demands. The inverse relationship between autophagy induction and primary cilia elongation offers mechanistic insight into how mTORC1 signaling may exert this control.

Previous work has implicated mTORC1 signaling in primary cilia biology, albeit with varied outcomes depending on the cellular context. For instance, studies in zebrafish and in cultured mammalian epithelial cells have reported opposing effects of mTORC1 inhibition on primary cilia length, with shortening in one context and elongation in another^33,34^. Similarly, genetic models of constitutive mTORC1 activation have shown divergent results: *Tsc1*-null MEFs exhibited elongated cilia reversible by rapamycin, while *Tsc2-*null MEF cells displayed shorter primary cilia and reduced overall ciliogenesis^36^. In contrast, it has also been reported that both *Tsc1* and *Tsc2* null MEFs exhibit increased ciliogenesis, with rapamycin further enhancing this effect, and increased cilia length, which appeared resistant to rapamycin^35^. A study of *Tsc2*-deficient mouse neurons found that they exhibit a rapamycin-reversible decrease in ciliation, with neuronal cilia length being unaffected by *Tsc2* knockdown^37^. These discrepant findings might highlight the complexity of the relationship between mTORC1 signaling and primary cilia in different settings and underscore a need for uniform experimental conditions in isogenic settings where effects on cell size and cell cycle can be assessed in parallel.

Several studies have previously identified links between autophagy and ciliogenesis, although with discrepant conclusions and few studies reporting effects on cilia length. Tang et al. demonstrated that autophagy promotes ciliogenesis by degrading OFD1 at centriolar satellites, and that disruption of this autophagic process impairs ciliary assembly^64^. In contrast, Pampliega et al. found that autophagy negatively regulates ciliogenesis by degrading IFT20 and other intraflagellar transport proteins involved in ciliary assembly, with autophagy inhibition resulting in enhanced ciliogenesis and, as also reported here, cilia elongation^65^. Neither of these studies systematically measured primary cilia length under controlled and monitored cell-cycle conditions, which are tightly linked to cilia formation and disassembly^27–30^. Since primary cilia are assembled during G0/G1 and disassembled upon cell-cycle re-entry, it is critical to account for cell-cycle state when assessing primary cilia formation or length in proliferating cultured cells. In contrast, in terminally differentiated primary cells or intact tissues where most cells are non-proliferative, cell cycle influence is likely minimal.

The study here controls for many of these experimental differences that can influence primary cilia dynamics. Through both pharmacological and genetic approaches and in distinct cell types and species, we demonstrate a consistent relationship between mTORC1 signaling and primary cilia length that is mechanistically linked to autophagy suppression but independent of effects on the cell cycle or cell size. Our results align with a previous study by Maharjan et al. that showed autophagy inhibition prevents cilia disassembly in RPE1 cells, whereas our findings demonstrate that the mTORC1-mediated suppression of autophagy promotes primary cilia lengthening^66^. Both studies show that autophagy inhibition preserves or elongates primary cilia, albeit under distinct experimental conditions, with this current study extending these findings to growth factor-independent mTORC1 signaling.

## LIMITATIONS OF STUDY

Future mechanistic studies detailing the regulation of mTORC1-autophagy–mediated control of primary cilia length are warranted. For instance, are the effects on primary cilia growth a component of bulk autophagy or are primary cilia components specifically targeted for autophagic degradation. It also remains unclear whether mTORC1 modulates specific ciliary signaling pathways such as those downstream of the Sonic Hedgehog or Wnt ligands, which have been found to be propagated via primary cilia^67–74^. Furthermore, although our data support a general mechanism functional in different cell types and species, it remains to be determined whether this regulation is conserved across all cell types, tissues, and physiological conditions where mTORC1 signaling may differentially respond.

These findings establish mTORC1 as a regulator of primary cilia length through suppression of autophagy, linking cellular metabolic status to one of the key organelles essential for intracellular communication. Thus, the primary cilia is an effector of mTORC1 signaling that should be considered for a better understanding of ciliopathies and as a potential consequence of dysregulated mTORC1 signaling.

## ACKNOWLEDGEMENTS

We thank Dr. Angelica D’Amore, Emma Martin, and Dr. Mustafa Sahin for technical advice on isolation and culture of cortical neurons from mouse embryos. We thank Eric Marino, Dr. Zhe Cao, and Dr. Christopher Morrow for technical advice on confocal microscopy. This work was supported by grants from the Glenn Foundation for Medical Research Postdoctoral Fellowship to Y.C., NIH (T32-DK128781) to S.C.L. and K.C.K., NIH (F31-DK128873) to K.C.K, Damon Runyon Cancer Research Foundation Merck Fellowship DRG-#2443-21 to M.Y.C., and NIH (R21-NS126952, R35-CA197459, P01-CA120964) to B.D.M.

## AUTHOR CONTRIBUTIONS

Conceptualization, J.S. and B.D.M.; investigation, J.S., P.N., S.S., K.T., Y.C., S.L.C., M.Y.C., K.C.K.; writing, J.S. and B.D.M.; funding acquisition, S.S., Y.C., S.L.C., M.Y.C., K.C.K., and B.D.M., resources, B.D.M., supervision, B.D.M.

## DECLARATION OF INTERESTS

All authors declare no competing financial interests.

## RESOURCE AVAILABILITY

### Lead contact

Further information and requests for resources and reagents should be directed to and will be fulfilled by the lead contact, Brendan Manning.

### Materials availability

Unique materials generated in this study are available from the Lead contact without restriction.

### Data and code availability

- All original data reported in this paper will be shared by the Lead contact upon request.
- This paper does not report original code.
- Any additional information required to reanalyze the data reported in this paper is available from the Lead contact upon request.

## MATERIALS AND METHODS

### KEY RESOURCES TABLE

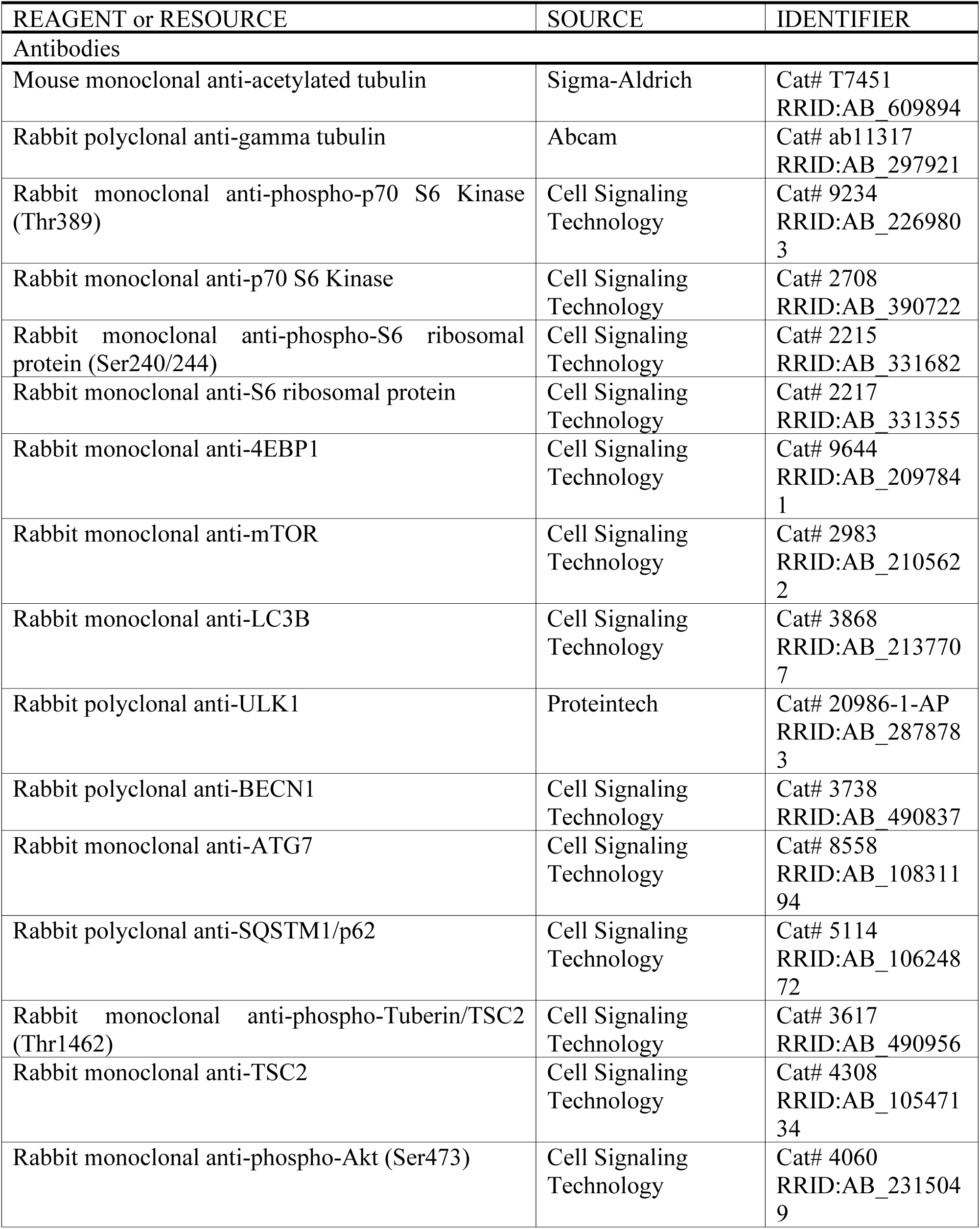
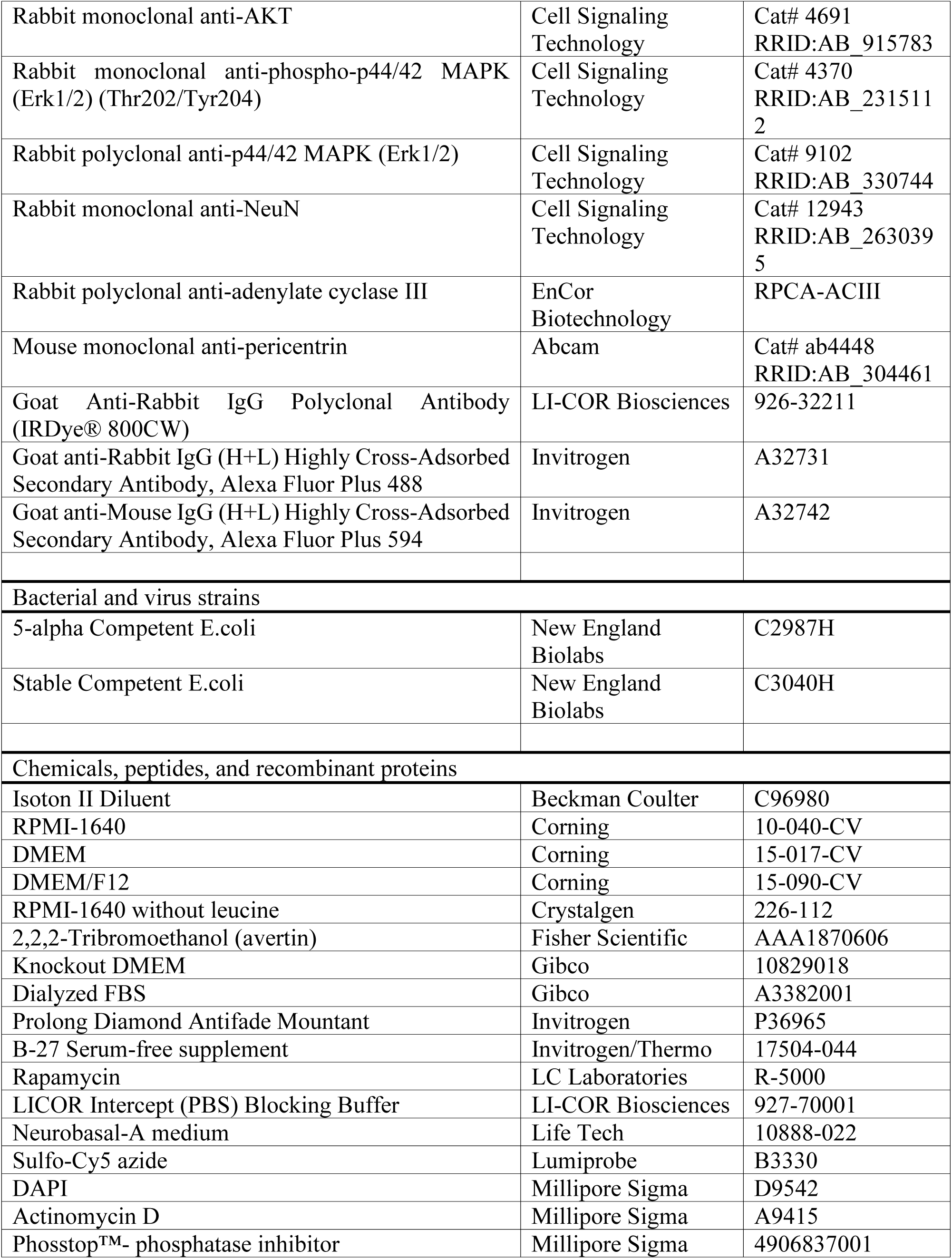
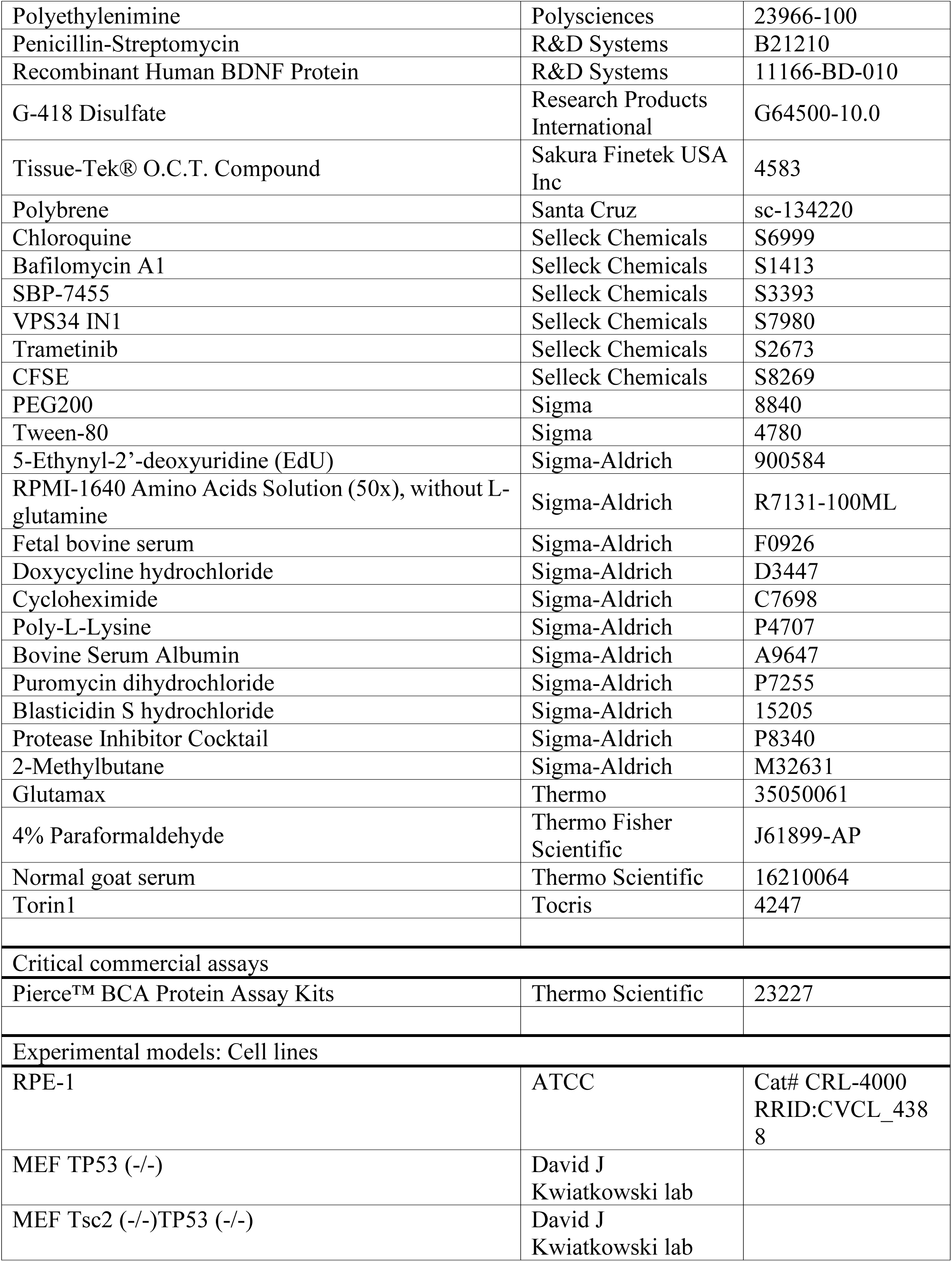
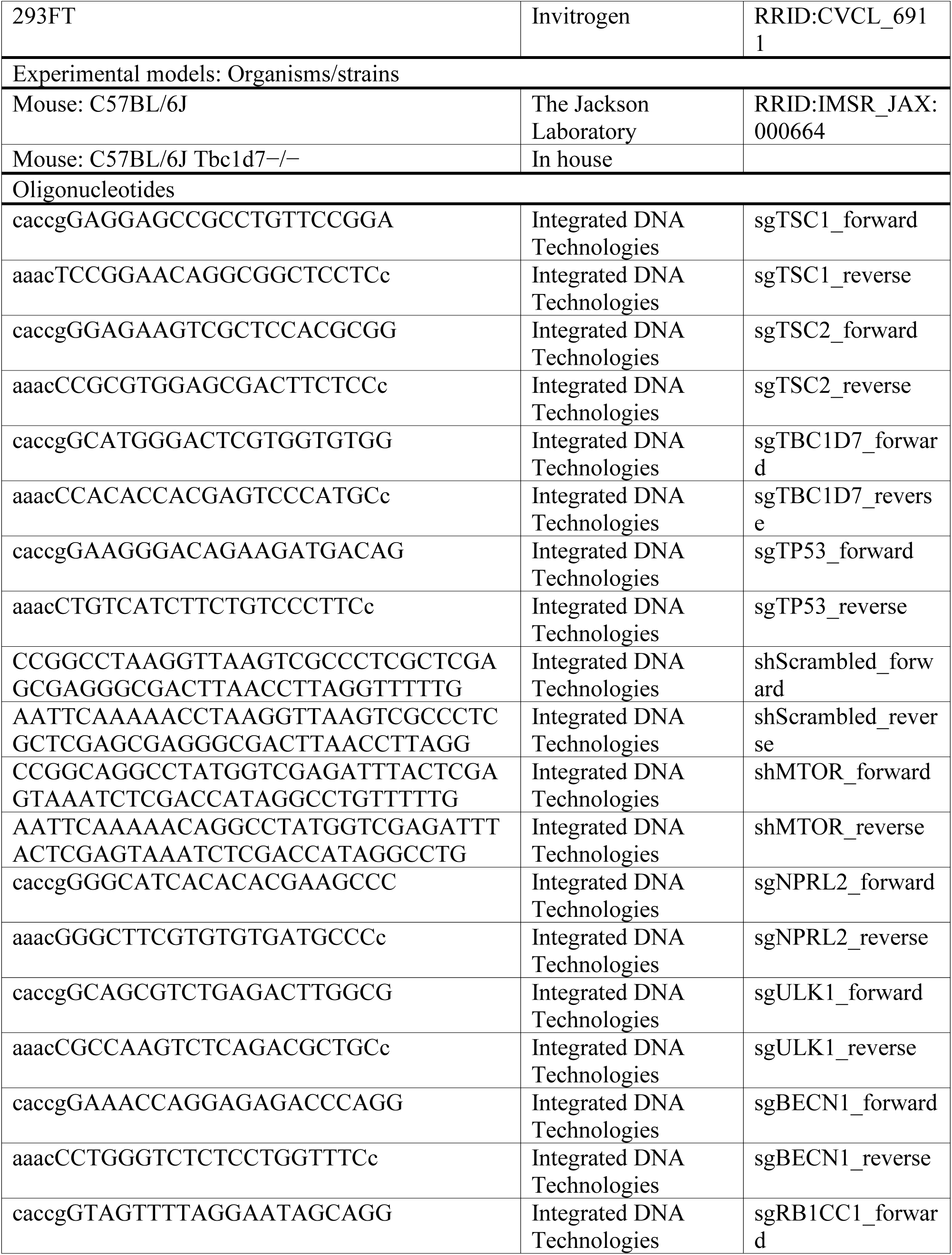
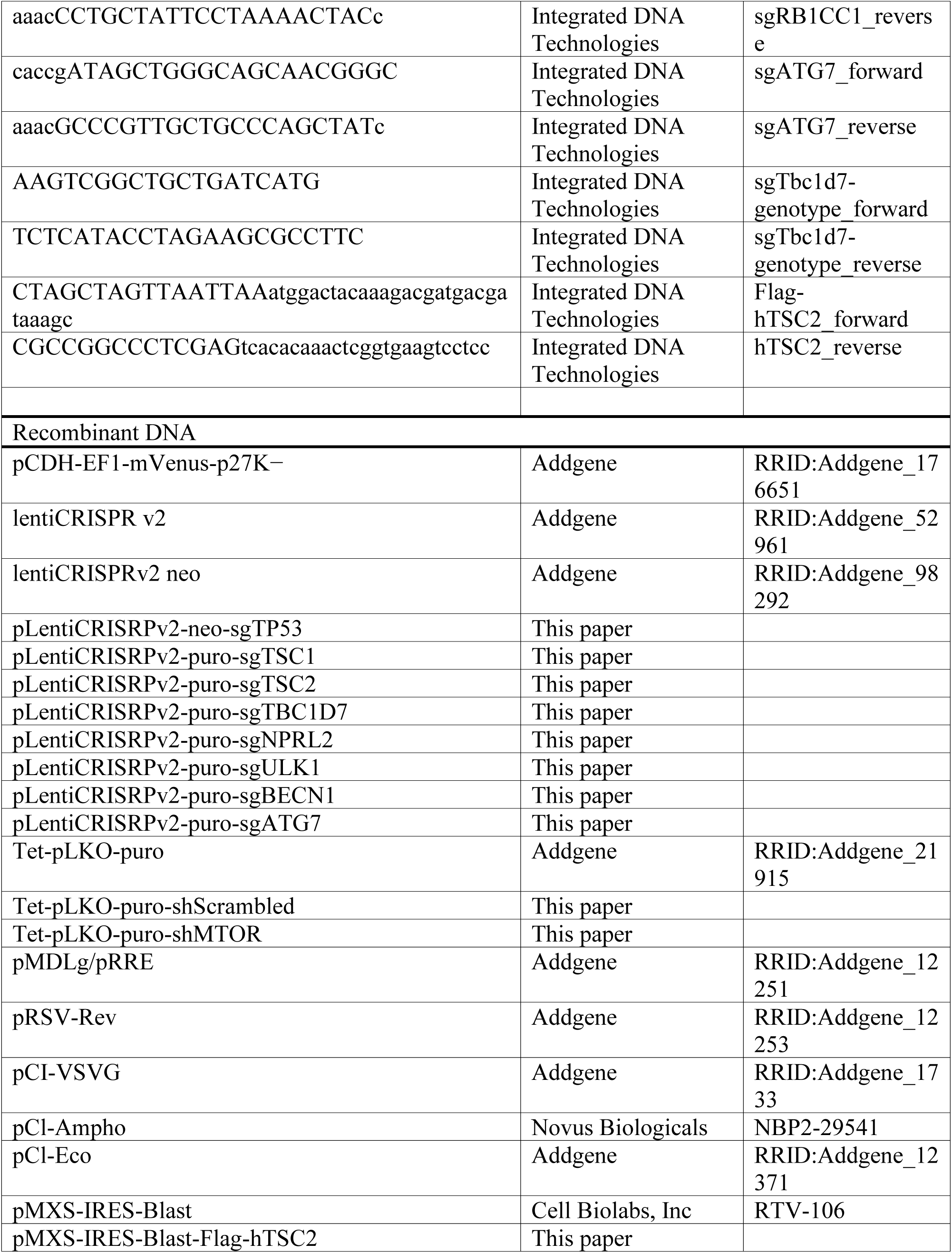
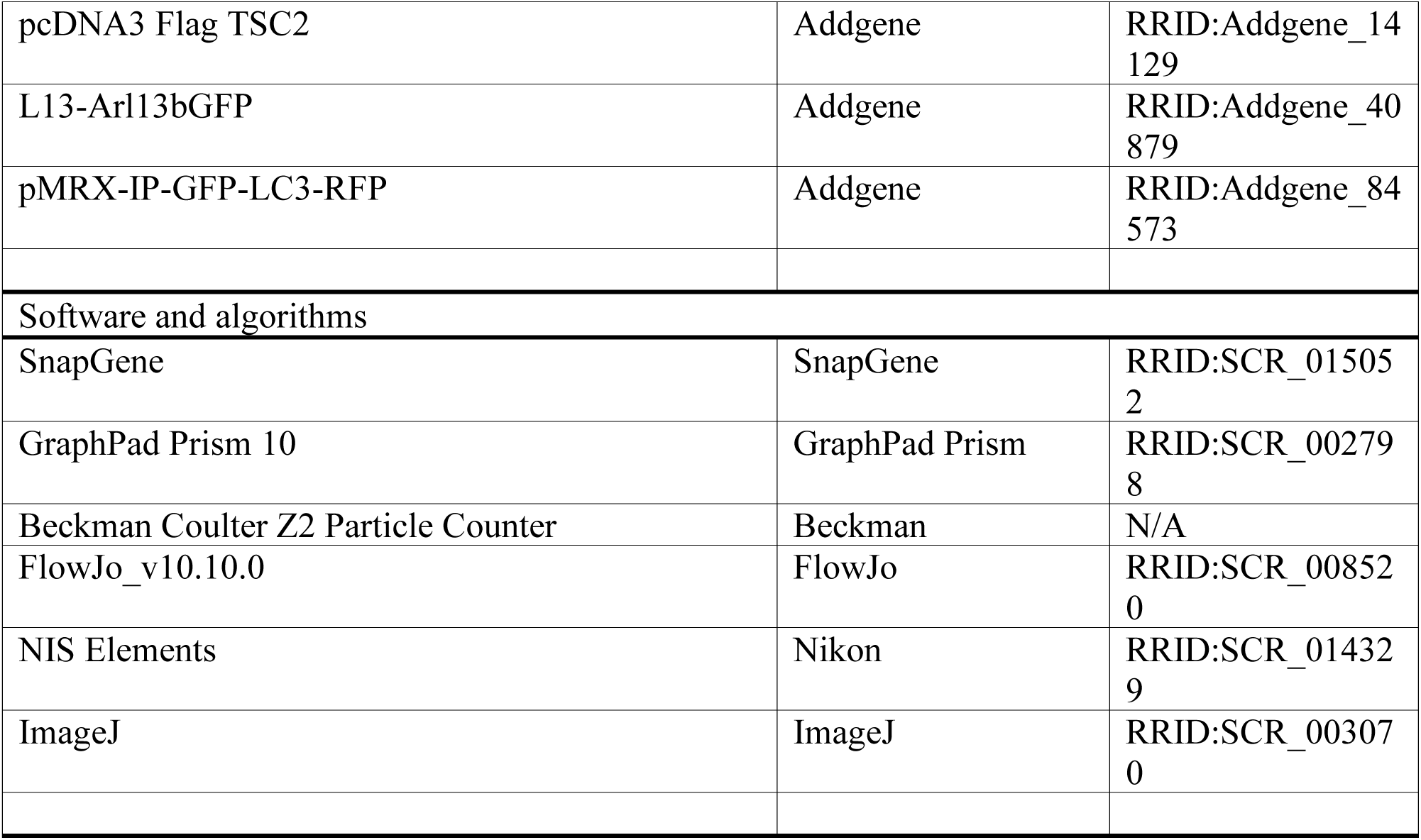

### EXPERIMENTAL MODEL AND STUDY PARTICIPANT DETAILS

#### Animals

All animal studies were reviewed and approved by the Harvard Medical Area Standing Committee on Animals IACUC, an AAALAC International- and USDA-accredited facility. All animal care and protocols adhered to the principles outlined in the Guide for the Care and Use of Laboratory Animals. Mice were group-housed (3–4 per cage with same-sex littermates after weaning at postnatal day 21) in temperature-controlled, pathogen-free facilities with a 12:12 h light-dark cycle in standard static microisolator-top cages. Autoclaved food (LabDiet; 5058) and water were provided ad libitum.

To prepare E16.5 embryos for cortical neuron isolation, three pairs of male and female C57BL/6J mice (Jackson Laboratory; 000664; 8–9 weeks old) or *Tbc1d7^+/−^* and *Tbc1d7^−/−^* mice (in-house; 8– 9 weeks old) were set up for timed pregnancies by separating the males from the females one day after breeding. Pregnancy progression was monitored over 1.5–2 weeks after separation. The pregnant females were then subjected to procedures for embryonic cortical neuron isolation.

To pharmacologically perturb mTORC1 signaling in vivo, rapamycin (1 mg/kg) was administered daily via intraperitoneal injection to 8–9-week-old male mice for one week. Male mice were selected to minimize hormonal variations that could introduce additional variables in mTORC1 signaling outcomes. Before administration, each mouse’s weight was measured and recorded. The appropriate volume of rapamycin solution, corresponding to the targeted dose, was calculated based on the recorded weight and injected into the subject. Control mice received an equivalent volume of ethanol, prepared in parallel with the rapamycin solution.

Rapamycin (LC Laboratories; R-5000) was resuspended in 100% ethanol at a concentration of 2.5 mg/mL and stored at −80°C in small aliquots to prevent repeated freeze-thaw cycles. On the day of treatment, the rapamycin stock was diluted to 0.125 mg/mL in a resuspension buffer prepared as follows: 45 mL sterile PBS, 2.5 mL PEG200 (Sigma; 8840), and 2.5 mL Tween-80 (Sigma; 4780). The resuspension buffer was filtered through a 0.2 µm sterile filter and stored at 4°C until use. Freshly diluted rapamycin was prepared for each experiment to ensure consistency and drug stability.

#### Cultured primary neurons

Using aseptic techniques, the uterine horn of freshly euthanized pregnant females containing E16.5 embryos was removed and transferred to a 10-cm tissue culture plate containing cold Neurobasal medium (Life Tech; 10888-022). Under a stereomicroscope (Olympus; SZX7), the head of each embryo was resected and opened to extract the brain. The hemispheres were separated, and the meninges were removed. The neocortex was dissected and transferred to a 6-cm tissue culture plate containing cold Neurobasal medium. All neocortices were pooled, except for those from embryos derived from crosses between *Tbc1d7^+/−^* and *Tbc1d7^−/−^*mice, which were processed separately to prevent cross-contamination between genotypes. Tail tissues were collected for subsequent genomic DNA extraction and PCR genotyping.

The collected neocortices were centrifuged at 1,000 × g for 3 minutes, and the supernatant was aspirated. The cortical tissues were mechanically dissociated by pipetting in cold Neurobasal medium and filtered through a 0.40 µm mesh filter. The cells were washed twice with cold Neurobasal medium, and the cell concentration was measured using a Beckman Coulter counter. Typically, two neocortices from an embryo yielded approximately 2 million cells. Approximately 300,000–500,000 cells/well or 30,000–50,000 cells/well were seeded in 12-well or 24-well plates, respectively, in Neurobasal A medium (Life Technologies; 10888-022) containing 2% B-27 (Invitrogen/Thermo; 17504-044), 1% GlutaMAX (Thermo; 35050061), 1% penicillin-streptomycin (R&D Systems; B21210), and 10 ng/mL BDNF (R&D Systems; 11166-BD-010). For immunofluorescence, cells were seeded onto 12 mm diameter coverslips (Fisher Scientific; 50-143-822) in 24-well plates. The coverslips were pre-washed (with distilled water, HCl, distilled water, and 100% ethanol) and coated with 0.01% poly-L-lysine (Sigma-Aldrich; P4707; 30 min incubation at room temperature). Neurons were cultured in a humidified incubator at 37°C with 5% CO₂. The medium was changed every 3–4 days. Neuronal sprouting was monitored until the day of harvest. All neuronal treatments were performed for one day starting at day 7 in vitro (DIV 7).

#### Cell lines

The hTERT-immortalized human retinal pigment epithelial (RPE1) cells (ATCC; CRL-4000) were cultured in RPMI-1640 supplemented with 10% FBS and 1% penicillin-streptomycin in a humidified incubator at 37°C with 5% CO₂. Mouse embryonic fibroblast (MEF) cell lines (*Tsc2^⁺/⁺^; TP53^⁻/⁻^* and *Tsc2^⁻/⁻^;TP53^⁻/⁻^,* from the lab of David J. Kwiatkowski)^17^ were cultured in DMEM supplemented with 10% FBS and 1% penicillin-streptomycin. HEK 293FT cells (Invitrogen; R70007) were cultured in KnockOut DMEM containing 10% FBS, 1% GlutaMAX, and 1% penicillin-streptomycin. All cells were maintained at 25–90% confluence and split when they became confluent using 0.05% trypsin with 0.53 mM EDTA (Corning; 25-052-Cl).

### Bacterial strains

To transform and clone plasmids, two commercial E. coli strains from New England Biolabs (NEB) were used: 5-alpha Competent E. coli (NEB; C2987H) was used for cloning retroviral vectors (e.g., pMXS), and Stable Competent E. coli (NEB; C3040H) was used for cloning lentiviral vectors (e.g., pLentiCRISPR).

### METHOD DETAILS

#### Cell culture

To induce the formation of primary cilia, cells were cultured in full growth medium (supplemented with 10% FBS and 1% penicillin-streptomycin) and then serum-starved by first washing with PBS once, followed by the addition of base medium without FBS supplementation for approximately two days. Drug treatments began one day after serum starvation and lasted for one day. For leucine or amino acid deprivation, the FBS-lacking medium was replaced at 24 hours post-serum starvation with RPMI-1640 lacking either leucine alone or all amino acids for a 24-hour period. To assess differential primary cilia elongation in various media, RPE1 cells were cultured in either RPMI-1640, DMEM (Corning; 15-017-CV), or DMEM/F-12 (Corning; 15-090-CV) supplemented with 10% FBS and 1% penicillin-streptomycin for one week to allow adaptation. Cells in these different media were then serum-starved with their respective FBS-free medium for 48 hours.

#### EdU incorporation for cell cycle analysis

Approximately 1 × 10⁶ adherent cells were incubated with 10 µM of 5-ethynyl-2’-deoxyuridine (EdU; Sigma-Aldrich; 900584) in culture medium for one hour. The cells were then trypsinized and fixed with cold 75% ethanol for as short as a few hours or up to one week at 4°C. The fixed cells were gently pelleted at 1,000 × g for two minutes (these parameters were maintained for all subsequent centrifugation steps) and washed once with phosphate-buffered saline (PBS) containing 5% bovine serum albumin (BSA; Sigma-Aldrich; A9647) and 2 mM ethylenediaminetetraacetic acid (EDTA). Cells were then permeabilized with 0.1% Triton X-100 (prepared in wash buffer) for 10 minutes at room temperature. After one wash, the Click-iT EdU labeling reaction was performed by incubating the cells in darkness for 30 minutes with the following freshly prepared labeling mixture: 175.8 µL PBS, 4 µL 100 mM copper sulfate pentahydrate (CuSO₄; freshly made), 0.2 µL 4 mM sulfo-Cy5 azide (Lumiprobe; B3330), and 20 µL 200 mg/mL ascorbate (freshly made). The Cy5-labeled cells were washed once and incubated 1 min with 1 µg/mL 4’,6-diamidino-2-phenylindole (DAPI; Millipore Sigma; D9542) for genomic DNA staining. Cy5- and DAPI-labeled cells were analyzed using the LSR Fortessa (Becton Dickinson) and FlowJo software (FlowJo).

#### CFSE labeling

For intracellular protein quantification, RPE1 cells were subjected to drug treatments as indicated, then washed twice with PBS and labeled with carboxyfluorescein succinimidyl ester (CFSE; final concentration 10 µM; ThermoFisher Scientific). CFSE was freshly diluted from a 10 mM stock in pre-warmed PBS. Cells were incubated with the CFSE labeling solution at 37°C for 1 hour in the dark. Following incubation, cells were washed twice with PBS, trypsinized, and resuspended in PBS for flow cytometric analysis. Fluorescence intensity, which reflects intracellular protein abundance due to CFSE’s covalent binding to amine groups on proteins, was measured in the FITC channel using a BD LSRFortessa analyzer^75^.

#### Cell proliferation and growth

Cell proliferation was measured using a Beckman Coulter Z2 particle counter (Beckman Coulter). Approximately 5 × 10⁴ cells were seeded in a 12-well plate overnight. The next day, the culture medium was replaced with serum-starvation medium (lacking FBS). At 24 hours post-serum starvation, FBS or drug (rapamycin or Torin1) was added. After 24 hours of treatment, cells were trypsinized and counted using the Beckman Coulter Z2 particle counter, measuring cell diameters between 8 and 30 µm. To prepare cell samples for counting, 200 µL of the trypsinized cells (from one biological replicate) was diluted in 9.8 mL of Isoton II Diluent (1:500 dilution; Beckman Coulter; C96980) and mixed in a 25 mL Accuvette cup (Beckman Coulter; A35473). The resulting count (cells/mL) was multiplied by 100 to obtain the total number of cells in the biological sample. Cell volume measurements were performed using the same sample preparation method.

#### Endogenous gene deletion using CRISPR-Cas9

Endogenous gene deletion was achieved using the CRISPR-Cas9 system, targeting specific gene loci with single-guide RNAs (sgRNAs). The CRISPR-Cas9 and sgRNA constructs were delivered via lentiviral transduction. The lentiviral plasmid lentiCRISPRv2 (pLentiCRISPRv2), a gift from Feng Zhang (puromycin selection; Addgene plasmid # 52961; http://n2t.net/addgene:52961)^76^, or for targeting p53 (neomycin selection, lentiCRISPRv2-neo; Addgene; 98292)^77^, was modified by inserting the following sgRNA sequences at the BsmB1 site, with inserts prepared by annealing two primers, each containing a flanking sequence recognizing the BsmB1-digested lentiCRISPRv2 plasmid (sgRNA sequences in uppercase, Bsmb1-annealing sequences in lowercase):

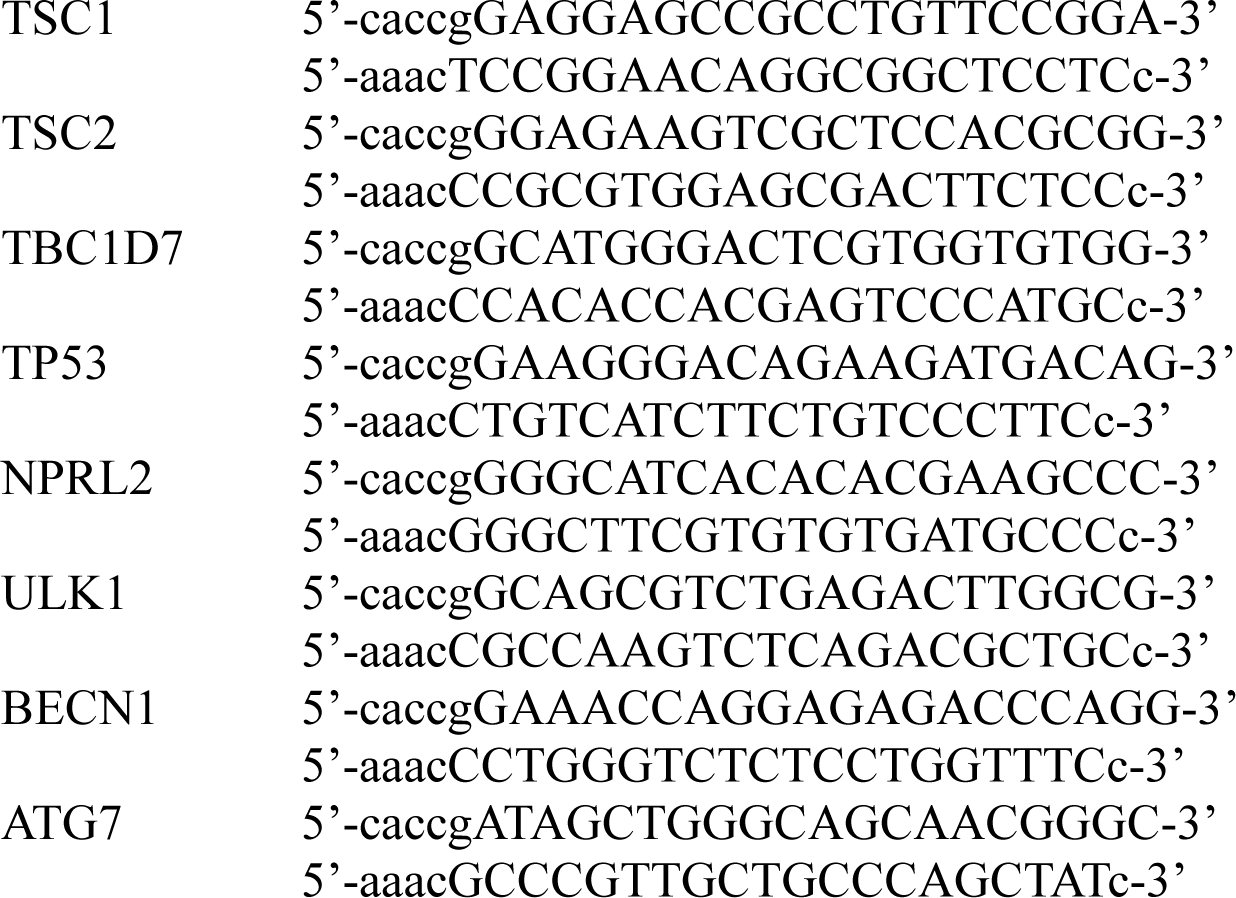

To anneal primers (Integrated DNA Technologies), 100 µM of each were incubated in 1X NEBuffer 2 (New England Biolabs [NEB]; B7002S) at 94°C for 2 min, followed by 65°C for 10 min and 4°C for 1 min. The annealed primers (diluted 1:100) were inserted into pLentiCRISPRv2 digested by the BsmB1 enzyme (NEB; R0739S). Ligation was performed using the Quick Ligation Reaction Kit (NEB; M2200S) according to the manufacturer’s instructions. The ligated plasmids were transformed into NEB Stable competent E. coli (NEB; C3040H) with ampicillin selection. Plasmids were purified using the QIAprep Spin Miniprep Kit (Qiagen; 27106) and correct sgRNA insertion was verified by whole plasmid sequencing using Oxford Nanopore Technology (Plasmidsaurus).

Pseudotyped lentivirus encoding pLentiCRISPRv2-sgRNA was produced by transfecting 293FT cells (Invitrogen; R70007) with pLentiCRISPRv2, pMDLg/pRRE, pRSV-Rev (third-generation lentiviral packaging plasmids), and pCl-VSVG (envelope plasmid). Transfection was performed using polyethylenimine (PEI; Polysciences; 23966-100) at a 1:3 DNA-to-PEI ratio. A total of 6 µg of PEI was mixed with 1 µg pLentiCRISPRv2-sgRNA, 0.5 µg pMDLg/pRRE, 0.5 µg pRSV-Rev, and 0.1 µg pCl-VSVG, and the PEI-DNA complex was used to transfect approximately 7–8 × 10⁵ 293FT cells in a 6 cm dish. The next day, the culture medium was changed, and cells were allowed to recover for 24 hours. After 24 hours post-media change, 1% BSA was added to enhance viral titers. Culture medium containing lentivirus was collected daily for up to 96 hours post-media change and filtered through a 0.45 µm pore-sized filter.

For lentiviral delievery, approximately 1 × 10⁵ wild-type target cells were seeded in a 6-well plate overnight. The next day, after changing the culture medium, the filtered lentivirus was added along with 8 µg/mL polybrene (Santa Cruz; sc-134220). The medium was changed the following day, and cells were allowed to recover. Selection of infected cells was performed with 10 µg/mL puromycin (Sigma-Aldrich; P7255) for 48–72 hours or 500 µg/mL G-418 (Research Products International; G64500-10.0) for one week.

To obtain single-cell clones (e.g., sgTSC1, sgTSC2, and sgTBC1D7 RPE1 cells), puromycin-selected cells were plated in 96-well plates at a single-cell density. Only wells containing a single parental-like cell were closely monitored, and upon reaching high cell density, clones were expanded into larger plate formats. During this process, cells were cultured with 5 µg/mL plasmocin (Invivogen; ant-mpt) to prevent mycoplasma contamination. Single-cell clones were validated by the absence of target protein expression and functional changes (e.g., growth factor-independent activation of mTORC1 signaling in sgTSC1, sgTSC2, or sgTBC1D7 cells).

#### Transduction of TSC2

Flag-tagged human TSC2 (Flag-TSC2) cDNA from pcDNA3 Flag TSC2 (Addgene; 14129)^18^ was PCR-amplified using Phusion High-Fidelity DNA Polymerase (Thermo Scientific; F531S) with primers containing sequences homologous to the insertion site in pMXS-IRES-Blast, a retroviral vector (Cell Biolabs, Inc; RTV-106):

5’-CTAGCTAGTTAATTAAatggactacaaagacgatgacgataaagc-3’

5’-CGCCGGCCCTCGAGtcacacaaactcggtgaagtcctcc-3’

(Uppercase: homologous to pMXS-IRES-Blast; lowercase: homologous to Flag-TSC2).

The PCR product was resolved by DNA electrophoresis, gel-extracted using the QIAquick Gel Extraction Kit (Qiagen; 28704), and inserted into pMXS-IRES-Blast linearized by PacI (NEB; R0547S) and XhoI (NEB; R0146S) using NEBuilder HiFi DNA Assembly (NEB; E2621S). The plasmid was cloned in NEB 5-alpha competent E. coli (NEB; C2987H). Successful Flag-TSC2 insertion was verified by whole plasmid sequencing (Plasmidsaurus).

Retroviral infection of sgTSC2 RPE1 cells was performed using pMXS-IRES-Blast-Flag-TSC2 and pCl-Ampho (a retroviral packaging vector), following the same transfection and infection protocols as described above for lentiviral-mediated gene transduction. Selection of transduced cells was performed using 10 µg/mL blasticidin (Sigma-Aldrich; 15205).

#### Generation of primary cilia and autophagy reporter cell lines

A primary cilia reporter cell line was generated by transducing GFP-tagged Arl13B into target RPE1 cells. Pseudotyped lentivirus carrying Arl13B-GFP was produced using the same method as described above for pLentiCRISPR lentivirus. The lentiviral plasmid expressing Arl13B-GFP (L13-Arl13bGFP) was a gift from Tamara Caspary (Addgene plasmid #40879; http://n2t.net/addgene:40879)^44^. Upon treatments (e.g., Torin1), live Arl13B-GFP RPE1 cells were seeded on a 35 mm Petri dish with a 14 mm glass-bottom microwell (MatTek; P35G-1.5-14-C). Primary cilia length was analyzed using an Inverted Nikon C2+ laser scanning confocal microscope.

Similarly, an autophagy reporter cell line was generated by transducing GFP-LC3-RFP into target RPE1 cells. Pseudotyped retrovirus carrying GFP-LC3-RFP was produced using the same method as described for Flag-TSC2 retrovirus. The retroviral plasmid expressing GFP-LC3-RFP (pMRX-IP-GFP-LC3-RFP) was a gift from Noboru Mizushima (Addgene plasmid #84573; http://n2t.net/addgene:84573)^78^. Fluorescently labeled cells were confirmed and further sorted using a BD Aria sorter, operated by the Flow Cytometry Core in the Department of Immunology at Harvard Medical School. Upon treatments (e.g., serum starvation or drug treatment), live GFP-LC3-RFP cells were analyzed for differential autophagy flux using an LSR Fortessa flow cytometer (Becton Dickinson).

#### Generation of Dox-inducible shMTOR cell line

A doxycycline-inducible mTOR shRNA-expressing RPE1 cell line (shMTOR) was generated using the Tet-pLKO-Puro vector system (a gift from Dmitri Wiederschain; Addgene plasmid #21915; http://n2t.net/addgene:21915)^79^. The shMTOR insert sequence used for cloning was:

5’-ccggCAGGCCTATGGTCGAGATTTActcgagTAAATCTCGACCATAGGCCTGtttttg-3’

5’-aattcaaaaaCAGGCCTATGGTCGAGATTTActcgagTAAATCTCGACCATAGGCCTG-3’

The Tet-pLKO-Puro plasmid was linearized by digestion with AgeI (NEB; R3552S) and EcoRI (NEB; R3101S) restriction enzymes to generate complementary overhangs for the annealed shMTOR oligonucleotides. A total of 2 µg of vector DNA was digested at 37°C for 3 hours. The digested vector was resolved by DNA gel electrophoresis and purified using the QIAquick Gel Extraction Kit (Qiagen; 28704) according to the manufacturer’s protocol. The shMTOR forward and reverse oligonucleotides were annealed to form a double-stranded insert. Equimolar concentrations (100 µM) of both oligonucleotides were mixed in 1X NEBuffer 2 (NEB; B7002S), heated at 94°C for 5 minutes, and then gradually cooled to room temperature for annealing. The annealed shMTOR duplex was ligated into the linearized Tet-pLKO-Puro plasmid using the Quick Ligation Reaction Kit (NEB; M2200S) according to the manufacturer’s instructions. The ligation product was then transformed into NEB Stable competent E. coli cells (NEB; C3040H), with ampicillin (100 µg/mL) selection. Plasmids from selected colonies were extracted using the QIAprep Spin Miniprep Kit (Qiagen; 27106) and verified by whole plasmid sequencing via Plasmidsaurus using Oxford Nanopore Technology.

Lentivirus encoding shMTOR was produced by co-transfecting 293FT cells (Invitrogen; R70007) with Tet-pLKO-shMTOR, pMDLg/pRRE, pRSV-Rev (third-generation lentiviral packaging plasmids), and pCl-VSVG (envelope plasmid) using polyethylenimine (PEI; Polysciences; 23966-100) at a 1:3 DNA-to-PEI ratio. Lentiviral supernatant was collected at 48, 72, and 96 hours post-transfection, filtered through a 0.45 µm pore-sized filter, and stored at −80°C. For cell transduction, target cells were seeded at 1 × 10⁵ cells per well in a 6-well plate and transduced with filtered Tet-pLKO-shMTOR lentivirus in the presence of 8 µg/mL polybrene (Santa Cruz; sc-134220). After 24 hours, the medium was replaced, and cells were allowed to recover. Selection of stable clones was performed using 10 µg/mL puromycin (Sigma-Aldrich; P7255) for 72 hours.

To confirm inducible knockdown of MTOR, cells were treated with 0.5 µg/mL doxycycline (Sigma-Aldrich; D9891) for 5 days. Knockdown efficiency was assessed by Western blotting, verifying successful doxycycline-dependent knockdown of MTOR and suppression of mTORC1 signaling.

#### Genomic DNA isolation and genotyping of Tbc1d7 mouse strain

Tails from embryos derived from crosses between *Tbc1d7^+/−^*and *Tbc1d7^−/−^* mice were used for genotyping, as previously described^59^. Briefly, tail samples were resuspended in approximately 500 µL of lysis buffer containing 100 mM Tris (pH 8.0), 0.8% SDS, 5 mM EDTA, 400 mM NaCl, and 200 µg/mL proteinase K (NEB; P8107S), followed by incubation at 56°C overnight. The next day, the lysate was vigorously mixed with an equal volume of isopropanol. Precipitated genomic DNA was pelleted by centrifugation at 14,800 × g for 5 minutes at 4°C. The supernatant was removed, and the pellet was carefully washed twice with cold 70% ethanol. The purified DNA pellet was resuspended in 10 mM Tris (pH 8.0) and sheared by passing through a 27G needle 5–

10 times. The concentration of extracted genomic DNA was measured using a NanoDrop One/OneC Microvolume UV-Vis Spectrophotometer (Thermo Scientific; ND-ONE-W). Approximately 100 ng of genomic DNA was used for PCR amplification with Phusion High-Fidelity DNA Polymerase (Thermo Scientific; F531S) and the following primers:

5’-AAGTCGGCTGCTGATCATG-3’

5’-TCTCATACCTAGAAGCGCCTTC-3’

A size difference in the PCR products distinguishes the *Tbc1d7* exon 3 deletion from the wild-type allele, as indicated in ^59^.

#### Protein extracts and immunoblotting

About 1-2 x 10^6^ cells were harvested in a lysis buffer containing 50 mM HEPES pH 7.5, 40 mM NaCl, 2mM EDTA, 1% NP-40 (IGEPAL CA-630; Sigma; I8896), protease inhibitors (Sigma-Aldrich; P8340), and phosphatase inhibitors (Millipore Sigma; 4906837001). The lysates were briefly vortexed and incubated on ice for a few minutes before centrifugation at 15,000 × g at 4°C for 10 minutes to pellet the insoluble fraction. The supernatant (soluble fraction) was collected into a new tube for protein quantification using the Bicinchoninic Acid (BCA) assay (Thermo Scientific; 23227) according to the manufacturer’s instructions. Approximately 10–20 µg of total protein was mixed with 1X Laemmli sample buffer (125 mM Tris-HCl pH 6.8, 4% SDS, 20% glycerol, 0.004% bromophenol blue, and 10% β-mercaptoethanol), volumetrically normalized with lysis buffer, and heated at 95°C for 5–10 minutes. Proteins were resolved using a 4–15% Criterion TGX Precast Midi Protein Gel (Bio-Rad; 567-1084) in Tris-glycine running buffer (25 mM Tris-base, 0.1% SDS, 250 mM glycine) at a constant 120 V (Bio-Rad PowerPac Basic; Bio-Rad; 44457) for 80 minutes, until the Precision Plus Protein All Blue Prestained Protein Standards (Bio-Rad; 161-0393) fully migrated through the gel. After electrophoresis, the gel was transferred onto a 0.2 µm nitrocellulose membrane (Bio-Rad; 162-0112) using transfer buffer (50 mM Tris-base, 48 mM glycine, final pH 8.3) at a constant 300 mA for 1 hour. Successful protein transfer was confirmed by staining the membrane with Ponceau S solution (Sigma-Aldrich; P7170) for 5 minutes. The membrane was then cut into sections at known molecular weight regions, guided by the protein ladder, for immunolabeling with different primary antibodies (see Key Resource Table).

Membrane sections were briefly washed with TBST buffer (25 mM Tris-Base, 137 mM NaCl, 2.7 mM KCl, 0.1% Tween-20, pH 7.4) to remove Ponceau stain. They were then blocked for 30 minutes at room temperature in either: 5% dry milk in TBST (for antibodies against non-phosphorylated proteins; Stop & Shop Instant Dry Milk Non-Fat), or 5% BSA in TBST (for antibodies against phosphorylated proteins; Sigma-Aldrich; A9647). After blocking, membranes were incubated overnight at 4°C with primary antibodies diluted 1:1000 (or according to the manufacturer’s recommendation) in 5% dry milk or BSA in TBST. The next day, membranes were washed three times with TBST (5 minutes per wash at room temperature), then incubated with secondary antibody (1:10,000 dilution in 5% dry milk or BSA in TBST; LI-COR Biosciences; 926-32211) for 1 hour at room temperature. Following secondary antibody incubation, membranes were washed three times with TBST (5 minutes per wash at room temperature) and visualized via a Li-Cor Odyssey CLx system.

#### Immunofluorescence of primary cilia in cultured cells

Approximately 30,000-50,000 cells were seeded onto 12 mm diameter coverslips (Fisher Scientific; 50-143-822) in 24-well plates. Prior to culture, the coverslips were pre-washed (with distilled water, 2N HCl, distilled water, and 100% ethanol) and coated with 0.01% poly-L-lysine (Sigma-Aldrich; P4707; 30 min incubation at room temperature). Upon completion of the indicated treatments the medium was aspirated, and wells were washed once with PBS. Cells were then fixed with cold 100% methanol at 4°C for 10 minutes, washed twice with PBS, then permeabilized with 0.1% Triton X-100 in PBS for 10 minutes at room temperature. After permeabilization, samples were washed twice with PBS and incubated with LI-COR Intercept (PBS) Blocking Buffer (LI-COR Biosciences; 927-70001) for 30 minutes on a gentle shaker at room temperature. Samples were then incubated with a primary antibody mixture for 1 hour at room temperature, prepared in LI-COR Intercept (PBS) Blocking Buffer: For RPE1 and MEF cultures, anti-acetylated tubulin (monoclonal mouse; Sigma-Aldrich; T7451; 1:1000) and anti-gamma tubulin (polyclonal rabbit; Abcam; ab11317; 1:1000); For primary neurons, anti-adenylate cyclase III (polyclonal rabbit; EnCor Biotechnology; RPCA-ACIII; 1:500) and anti-pericentrin (monoclonal mouse; Abcam; ab4448; 1:1000). After primary antibody incubation, samples were washed three times with PBS and incubated with a secondary antibody mixture for 1 hour at room temperature, prepared in LI-COR Intercept (PBS) Blocking Buffer: anti-mouse Alexa Fluor Plus 594 (Invitrogen; A32742) and anti-rabbit Alexa Fluor Plus 488 (Invitrogen; A32731). Following incubation, the antibody mixture was removed, and samples were washed twice with PBS. Nuclei were stained by incubating samples with 1 µg/mL DAPI in PBS for 1 min. Coverslips were carefully lifted using fine-tip microforceps, then inverted onto a drop of ProLong Diamond Antifade Mountant (Invitrogen; P36965) on a clean Superfrost Plus Microscope Slide (Fisher Scientific; 12-550-15). Once slightly dried, coverslip edges were sealed with a thin layer of nail polish (VWR; 100491-940). After the nail polish dried, slides were stored in a polypropylene microscope slide box (Heathrow Scientific; 15991C) at 4°C until imaging with an inverted Nikon C2+ laser scanning confocal microscope.

#### Immunofluorescence of Primary Cilia in Mouse Brain Sections

Anesthetized vehicle and rapamycin-treated mice were subjected to a midline thoracic incision to expose the heart for transcardiac perfusion with 4% paraformaldehyde (PFA). A 25-gauge needle attached to a catheter tube was inserted into the left ventricle for perfusion. A small incision was made in the right atrium to allow outflow. Using a peristaltic pump (Ismatec Ism834C Reglo Digital Variable-Speed Peristaltic Pump), the circulatory system was flushed with PBS at a rate of ∼5 mL/min for 5 minutes. The perfusion was then switched to 4% PFA, continuing for 5–10 min, or until tissue stiffening and color change were observed.

Intact brains were extracted and immersed in 4% PFA for post-fixation at 4°C for 24 hours. The next day, 4% PFA was removed, and the brains were transferred to 15% sucrose (in PBS) at 4°C for 24 hours. The next day, 15% sucrose was replaced with 30% sucrose (in PBS) at 4°C until ready for cryosectioning. 2-methylbutane (Sigma-Aldrich; M32631) was poured into a small plastic box on dry ice to create a freezing environment, and OCT-embedding medium (Sakura Finetek USA Inc; 4583) was added to a cubical mold (Sigma-Aldrich; E6032) placed inside the box. The fixed brain was carefully oriented with the olfactory bulbs facing the bottom of the mold. The mold was left undisturbed until the OCT was completely frozen. Using a Microm cryostat (HM 505 N), the frozen brains were sectioned into 30-µm slices, which were collected in 6-well plates containing PBS. Brain slices were then washed three times with PBS (5 minutes per wash at room temperature) and permeabilized with 1% Triton X-100 in PBS (PBST) with gentle agitation overnight at 4°C. The next day, brain slices were incubated with 10% normal goat serum (NGS) in PBST for 1 hour at room temperature for blocking. Brain slices were incubated overnight at 4°C with primary antibodies diluted in 1% normal goat serum (PBST): anti-adenylate cyclase III (ACIII; EnCor Biotechnology; RPCA-ACIIIl 1:500) and anti-NeuN (Cell Signaling Technology; 12943; 1:1000). The next day, brain slices were washed three times with PBS (5 min each at room temperature) and incubated with a secondary antibody mixture for 2 hours at room temperature in 1% normal goat serum (PBST): anti-rabbit Alexa Fluor Plus 488 (Invitrogen; A32731; 1:1000) and anti-mouse Alexa Fluor Plus 594 (Invitrogen; A32742; 1:1000). Plates containing the specimens were covered with aluminum foil to minimize light exposure. Brain slices were washed three times with PBS (5 minutes per wash at room temperature) and incubated with 1 µg/mL DAPI in PBS for 2 minutes. Using a disposable transfer pipette (ULINE; S-24319), fully labeled brain slices were transferred onto clean Superfrost Plus Microscope Slides. Using a fine paintbrush, floating brain slices were carefully positioned, unfolded, and oriented properly on the microscope slide. Excess liquid was gently removed using a transfer pipette. Drops of ProLong Diamond Antifade Mountant were applied to each section, followed by the placement of coverslips and sealing with nail polish. After drying, slides were stored at 4°C until imaging using a Nikon C2+ laser scanning confocal microscope.

This immunostaining protocol was adapted in part from procedures described by EnCor Biotechnology [EnCor Biotechnology, “Procedure for immunofluorescent staining of free-floating brain sections,” EnCor Protocols, https://encorbio.com/protocols/immunots.htm].

#### Primary cilia length measurement

Lengths of primary cilia in cultured cells were measured by tracing the axoneme labeled with either acetylated tubulin or adenylate cyclase III, using the “Segmented Line” tool in ImageJ or the “Polyline” tool in Nikon Imaging Software (NIS) Elements. For live-cell imaging, primary cilia lengths in Arl13B-GFP-expressing RPE1 cells were measured by tracing the GFP-labeled axoneme at specific time points, such as pre-, during, and post-Torin1 treatment, using the “Polyline” tool in NIS Elements.

For the length measurement of primary cilia in brain tissue sections, confocal Z-stack images were acquired across a 30 µm Z-axis range with 0.3 µm step intervals. To facilitate accurate length measurement, the 3D-Z-stacked images of the region of interest were transformed into a 2D projection using the 3D Volume Viewer plugin in ImageJ, allowing all primary cilia within the field to be visualized on the same plane. The resulting 2D-rendered images were then analyzed using the “Segmented Line” tool in ImageJ to measure primary cilia length.

#### Quantification and statistical analysis

Statistical analysis was performed using GraphPad Prism 10 software. All data are presented as mean ± standard deviation (SD), as indicated. Outliers were identified and removed using the ROUT method with an aggressiveness Q of 1%. Comparisons between two groups were analyzed using an unpaired two-tailed Student’s t-test. Comparisons between three or more groups were performed using a two-way ANOVA followed by Šídák’s multiple comparisons test. The significance threshold (α) was set at 0.05, and all p-values are reported.

## Notes

### Competing Interest Statement

The authors have declared no competing interest.

## REFERENCES

1. Liu, G.Y., and Sabatini, D.M. (2020). mTOR at the nexus of nutrition, growth, ageing and disease. Nat Rev Mol Cell Biol 21, 183–203. 10.1038/s41580-019-0199-y.

2. Valvezan, A.J., and Manning, B.D. (2019). Molecular logic of mTORC1 signalling as a metabolic rheostat. Nat Metab 1, 321–333. 10.1038/s42255-019-0038-7.

3. Wolfson, R.L., and Sabatini, D.M. (2017). The Dawn of the Age of Amino Acid Sensors for the mTORC1 Pathway. Cell Metab 26, 301–309. 10.1016/j.cmet.2017.07.001.

4. Sekiguchi, T., Hirose, E., Nakashima, N., Ii, M., and Nishimoto, T. (2001). Novel G proteins, Rag C and Rag D, interact with GTP-binding proteins, Rag A and Rag B. J Biol Chem 276, 7246–7257. 10.1074/jbc.M004389200.

5. Sancak, Y., Peterson, T.R., Shaul, Y.D., Lindquist, R.A., Thoreen, C.C., Bar-Peled, L., and Sabatini, D.M. (2008). The Rag GTPases bind raptor and mediate amino acid signaling to mTORC1. Science 320, 1496–1501.

6. Bar-Peled, L., Chantranupong, L., Cherniack, A.D., Chen, W.W., Ottina, K.A., Grabiner, B.C., Spear, E.D., Carter, S.L., Meyerson, M., and Sabatini, D.M. (2013). A Tumor suppressor complex with GAP activity for the Rag GTPases that signal amino acid sufficiency to mTORC1. Science 340, 1100–1106. 10.1126/science.1232044340/6136/1100 [pii].

7. Panchaud, N., Peli-Gulli, M.P., and De Virgilio, C. (2013). Amino acid deprivation inhibits TORC1 through a GTPase-activating protein complex for the Rag family GTPase Gtr1. Sci Signal 6, ra42. 10.1126/scisignal.2004112.

8. Yang, H., Jiang, X., Li, B., Yang, H.J., Miller, M., Yang, A., Dhar, A., and Pavletich, N.P. (2017). Mechanisms of mTORC1 activation by RHEB and inhibition by PRAS40. Nature 552, 368–373. 10.1038/nature25023.

9. Dibble, C.C., Elis, W., Menon, S., Qin, W., Klekota, J., Asara, J.M., Finan, P.M., Kwiatkowski, D.J., Murphy, L.O., and Manning, B.D. (2012). TBC1D7 is a third subunit of the TSC1-TSC2 complex upstream of mTORC1. Mol Cell 47, 535–546. 10.1016/j.molcel.2012.06.009 S1097-2765(12)00504-7 [pii].

10. Inoki, K., Li, Y., Xu, T., and Guan, K.L. (2003). Rheb GTPase is a direct target of TSC2 GAP activity and regulates mTOR signaling. Genes Dev 17, 1829–1834. 10.1101/gad.1110003.

11. Stocker, H., Radimerski, T., Schindelholz, B., Wittwer, F., Belawat, P., Daram, P., Breuer, S., Thomas, G., and Hafen, E. (2003). Rheb is an essential regulator of S6K in controlling cell growth in Drosophila. Nat Cell Biol 5, 559–565. 10.1038/ncb995.

12. Saucedo, L.J., Gao, X., Chiarelli, D.A., Li, L., Pan, D., and Edgar, B.A. (2003). Rheb promotes cell growth as a component of the insulin/TOR signalling network. Nat Cell Biol 5, 566–571. 10.1038/ncb996.

13. Zhang, Y., Gao, X., Saucedo, L.J., Ru, B., Edgar, B.A., and Pan, D. (2003). Rheb is a direct target of the tuberous sclerosis tumour suppressor proteins. Nat Cell Biol 5, 578–581. 10.1038/ncb999.

14. Tee, A.R., Manning, B.D., Roux, P.P., Cantley, L.C., and Blenis, J. (2003). Tuberous sclerosis complex gene products, Tuberin and Hamartin, control mTOR signaling by acting as a GTPase-activating protein complex toward Rheb. Curr Biol 13, 1259–1268. 10.1016/s0960-9822(03)00506-2.

15. Garami, A., Zwartkruis, F.J., Nobukuni, T., Joaquin, M., Roccio, M., Stocker, H., Kozma, S.C., Hafen, E., Bos, J.L., and Thomas, G. (2003). Insulin activation of Rheb, a mediator of mTOR/S6K/4E-BP signaling, is inhibited by TSC1 and 2. Mol Cell 11, 1457–1466. 10.1016/s1097-2765(03)00220-x.

16. Kwiatkowski, D.J., Zhang, H., Bandura, J.L., Heiberger, K.M., Glogauer, M., el-Hashemite, N., and Onda, H. (2002). A mouse model of TSC1 reveals sex-dependent lethality from liver hemangiomas, and up-regulation of p70S6 kinase activity in Tsc1 null cells. Hum Mol Genet 11, 525–534. 10.1093/hmg/11.5.525.

17. Zhang, H., Cicchetti, G., Onda, H., Koon, H.B., Asrican, K., Bajraszewski, N., Vazquez, F., Carpenter, C.L., and Kwiatkowski, D.J. (2003). Loss of Tsc1/Tsc2 activates mTOR and disrupts PI3K-Akt signaling through downregulation of PDGFR. J Clin Invest 112, 1223–1233. 10.1172/JCI17222.

18. Manning, B.D., Tee, A.R., Logsdon, M.N., Blenis, J., and Cantley, L.C. (2002). Identification of the tuberous sclerosis complex-2 tumor suppressor gene product tuberin as a target of the phosphoinositide 3-kinase/akt pathway. Mol Cell 10, 151–162. 10.1016/s1097-2765(02)00568-3.

19. Inoki, K., Li, Y., Zhu, T., Wu, J., and Guan, K.L. (2002). TSC2 is phosphorylated and inhibited by Akt and suppresses mTOR signalling. Nat Cell Biol 4, 648–657. 10.1038/ncb839.

20. Menon, S., Dibble, C.C., Talbott, G., Hoxhaj, G., Valvezan, A.J., Takahashi, H., Cantley, L.C., and Manning, B.D. (2014). Spatial control of the TSC complex integrates insulin and nutrient regulation of mTORC1 at the lysosome. Cell 156, 771–785. 10.1016/j.cell.2013.11.049.

21. Ma, L., Chen, Z., Erdjument-Bromage, H., Tempst, P., and Pandolfi, P.P. (2005). Phosphorylation and functional inactivation of TSC2 by Erk implications for tuberous sclerosis and cancer pathogenesis. Cell 121, 179–193. 10.1016/j.cell.2005.02.031.

22. Crino, P.B., Nathanson, K.L., and Henske, E.P. (2006). The tuberous sclerosis complex. N Engl J Med 355, 1345–1356. 10.1056/NEJMra055323.

23. Kimmelman, A.C., and White, E. (2017). Autophagy and Tumor Metabolism. Cell Metab 25, 1037–1043. 10.1016/j.cmet.2017.04.004.

24. Breslow, D.K., and Holland, A.J. (2019). Mechanism and Regulation of Centriole and Cilium Biogenesis. Annu Rev Biochem 88, 691–724. 10.1146/annurev-biochem-013118-111153.

25. Sanchez, I., and Dynlacht, B.D. (2016). Cilium assembly and disassembly. Nat Cell Biol 18, 711–717. 10.1038/ncb3370.

26. Mill, P., Christensen, S.T., and Pedersen, L.B. (2023). Primary cilia as dynamic and diverse signalling hubs in development and disease. Nat Rev Genet 24, 421–441. 10.1038/s41576-023-00587-9.

27. Archer, F.L., and Wheatley, D.N. (1971). Cilia in cell-cultured fibroblasts. II. Incidence in mitotic and post-mitotic BHK 21-C13 fibroblasts. J Anat 109, 277–292.

28. Kasahara, K., and Inagaki, M. (2021). Primary ciliary signaling: links with the cell cycle. Trends Cell Biol 31, 954–964. 10.1016/j.tcb.2021.07.009.

29. Tucker, R.W., Pardee, A.B., and Fujiwara, K. (1979). Centriole ciliation is related to quiescence and DNA synthesis in 3T3 cells. Cell 17, 527–535. 10.1016/0092-8674(79)90261-7.

30. Tucker, R.W., Scher, C.D., and Stiles, C.D. (1979). Centriole deciliation associated with the early response of 3T3 cells to growth factors but not to SV40. Cell 18, 1065–1072. 10.1016/0092-8674(79)90219-8.

31. Brooks, R.F. (1976). Regulation of fibroblast cell cycle by serum. Nature 260, 248–250. 10.1038/260248a0.

32. Moreno-Cruz, P., Corral Nieto, Y., Manrique Garcia, L., Pereira, A.G., and Bravo-San Pedro, J.M. (2023). Protocols to induce and study ciliogenesis. Methods Cell Biol 175, 1–15. 10.1016/bs.mcb.2022.10.002.

33. Yuan, S., Li, J., Diener, D.R., Choma, M.A., Rosenbaum, J.L., and Sun, Z. (2012). Target-of-rapamycin complex 1 (Torc1) signaling modulates cilia size and function through protein synthesis regulation. Proc Natl Acad Sci U S A 109, 2021–2026. 10.1073/pnas.1112834109.

34. Sherpa, R.T., Atkinson, K.F., Ferreira, V.P., and Nauli, S.M. (2016). Rapamycin Increases Length and Mechanosensory Function of Primary Cilia in Renal Epithelial and Vascular Endothelial Cells. Int Educ Res J 2, 91–97.

35. Hartman, T.R., Liu, D., Zilfou, J.T., Robb, V., Morrison, T., Watnick, T., and Henske, E.P. (2009). The tuberous sclerosis proteins regulate formation of the primary cilium via a rapamycin-insensitive and polycystin 1-independent pathway. Hum Mol Genet 18, 151–163. 10.1093/hmg/ddn325.

36. Rosengren, T., Larsen, L.J., Pedersen, L.B., Christensen, S.T., and Moller, L.B. (2018). TSC1 and TSC2 regulate cilia length and canonical Hedgehog signaling via different mechanisms. Cell Mol Life Sci 75, 2663–2680. 10.1007/s00018-018-2761-8.

37. Di Nardo, A., Lenoel, I., Winden, K.D., Ruhmkorf, A., Modi, M.E., Barrett, L., Ercan-Herbst, E., Venugopal, P., Behne, R., Lopes, C.A.M., et al. (2020). Phenotypic Screen with TSC-Deficient Neurons Reveals Heat-Shock Machinery as a Druggable Pathway for mTORC1 and Reduced Cilia. Cell Rep 31, 107780. 10.1016/j.celrep.2020.107780.

38. Piperno, G., and Fuller, M.T. (1985). Monoclonal antibodies specific for an acetylated form of alpha-tubulin recognize the antigen in cilia and flagella from a variety of organisms. J Cell Biol 101, 2085–2094. 10.1083/jcb.101.6.2085.

39. Muresan, V., Joshi, H.C., and Besharse, J.C. (1993). Gamma-tubulin in differentiated cell types: localization in the vicinity of basal bodies in retinal photoreceptors and ciliated epithelia. J Cell Sci 104 (Pt 4), 1229–1237. 10.1242/jcs.104.4.1229.

40. Oki, T., Nishimura, K., Kitaura, J., Togami, K., Maehara, A., Izawa, K., Sakaue-Sawano, A., Niida, A., Miyano, S., Aburatani, H., et al. (2014). A novel cell-cycle-indicator, mVenus-p27K-, identifies quiescent cells and visualizes G0-G1 transition. Sci Rep 4, 4012. 10.1038/srep04012.

41. Thoreen, C.C., Kang, S.A., Chang, J.W., Liu, Q., Zhang, J., Gao, Y., Reichling, L.J., Sim, T., Sabatini, D.M., and Gray, N.S. (2009). An ATP-competitive mammalian target of rapamycin inhibitor reveals rapamycin-resistant functions of mTORC1. J Biol Chem 284, 8023–8032. 10.1074/jbc.M900301200.

42. Fingar, D.C., Salama, S., Tsou, C., Harlow, E., and Blenis, J. (2002). Mammalian cell size is controlled by mTOR and its downstream targets S6K1 and 4EBP1/eIF4E. Genes Dev 16, 1472–1487. 10.1101/gad.995802.

43. Nanda, J.S., and Lorsch, J.R. (2014). Labeling a protein with fluorophores using NHS ester derivitization. Methods Enzymol 536, 87–94. 10.1016/B978-0-12-420070-8.00008-8.

44. Larkins, C.E., Aviles, G.D., East, M.P., Kahn, R.A., and Caspary, T. (2011). Arl13b regulates ciliogenesis and the dynamic localization of Shh signaling proteins. Mol Biol Cell 22, 4694–4703. 10.1091/mbc.E10-12-0994.

45. Caspary, T., Larkins, C.E., and Anderson, K.V. (2007). The graded response to Sonic Hedgehog depends on cilia architecture. Dev Cell 12, 767–778. 10.1016/j.devcel.2007.03.004.

46. Shen, K., Valenstein, M.L., Gu, X., and Sabatini, D.M. (2019). Arg-78 of Nprl2 catalyzes GATOR1-stimulated GTP hydrolysis by the Rag GTPases. J Biol Chem 294, 2970–2975. 10.1074/jbc.AC119.007382.

47. Morita, K., Hama, Y., Izume, T., Tamura, N., Ueno, T., Yamashita, Y., Sakamaki, Y., Mimura, K., Morishita, H., Shihoya, W., et al. (2018). Genome-wide CRISPR screen identifies TMEM41B as a gene required for autophagosome formation. J Cell Biol 217, 3817–3828. 10.1083/jcb.201804132.

48. Yamamoto, A., Tagawa, Y., Yoshimori, T., Moriyama, Y., Masaki, R., and Tashiro, Y. (1998). Bafilomycin A1 prevents maturation of autophagic vacuoles by inhibiting fusion between autophagosomes and lysosomes in rat hepatoma cell line, H-4-II-E cells. Cell Struct Funct 23, 33–42. 10.1247/csf.23.33.

49. Mauthe, M., Orhon, I., Rocchi, C., Zhou, X., Luhr, M., Hijlkema, K.J., Coppes, R.P., Engedal, N., Mari, M., and Reggiori, F. (2018). Chloroquine inhibits autophagic flux by decreasing autophagosome-lysosome fusion. Autophagy 14, 1435–1455. 10.1080/15548627.2018.1474314.

50. Egan, D., Kim, J., Shaw, R.J., and Guan, K.L. (2011). The autophagy initiating kinase ULK1 is regulated via opposing phosphorylation by AMPK and mTOR. Autophagy 7, 643–644. 10.4161/auto.7.6.15123.

51. Kim, J., Kundu, M., Viollet, B., and Guan, K.L. (2011). AMPK and mTOR regulate autophagy through direct phosphorylation of Ulk1. Nat Cell Biol 13, 132–141. 10.1038/ncb2152.

52. Kamada, Y., Funakoshi, T., Shintani, T., Nagano, K., Ohsumi, M., and Ohsumi, Y. (2000). Tor-mediated induction of autophagy via an Apg1 protein kinase complex. J Cell Biol 150, 1507–1513. 10.1083/jcb.150.6.1507.

53. Ren, H., Bakas, N.A., Vamos, M., Chaikuad, A., Limpert, A.S., Wimer, C.D., Brun, S.N., Lambert, L.J., Tautz, L., Celeridad, M., et al. (2020). Design, Synthesis, and Characterization of an Orally Active Dual-Specific ULK1/2 Autophagy Inhibitor that Synergizes with the PARP Inhibitor Olaparib for the Treatment of Triple-Negative Breast Cancer. J Med Chem 63, 14609–14625. 10.1021/acs.jmedchem.0c00873.

54. Kihara, A., Noda, T., Ishihara, N., and Ohsumi, Y. (2001). Two distinct Vps34 phosphatidylinositol 3-kinase complexes function in autophagy and carboxypeptidase Y sorting in Saccharomyces cerevisiae. J Cell Biol 152, 519–530. 10.1083/jcb.152.3.519.

55. Bago, R., Malik, N., Munson, M.J., Prescott, A.R., Davies, P., Sommer, E., Shpiro, N., Ward, R., Cross, D., Ganley, I.G., and Alessi, D.R. (2014). Characterization of VPS34-IN1, a selective inhibitor of Vps34, reveals that the phosphatidylinositol 3-phosphate-binding SGK3 protein kinase is a downstream target of class III phosphoinositide 3-kinase. Biochem J 463, 413–427. 10.1042/BJ20140889.

56. Komatsu, M., Waguri, S., Koike, M., Sou, Y.S., Ueno, T., Hara, T., Mizushima, N., Iwata, J., Ezaki, J., Murata, S., et al. (2007). Homeostatic levels of p62 control cytoplasmic inclusion body formation in autophagy-deficient mice. Cell 131, 1149–1163. 10.1016/j.cell.2007.10.035.

57. Fuchs, J.L., and Schwark, H.D. (2004). Neuronal primary cilia: a review. Cell Biol Int 28, 111–118. 10.1016/j.cellbi.2003.11.008.

58. Li, X., Yang, S., Deepak, V., Chinipardaz, Z., and Yang, S. (2021). Identification of Cilia in Different Mouse Tissues. Cells 10. 10.3390/cells10071623.

59. Schrotter, S., Yuskaitis, C.J., MacArthur, M.R., Mitchell, S.J., Hosios, A.M., Osipovich, M., Torrence, M.E., Mitchell, J.R., Hoxhaj, G., Sahin, M., and Manning, B.D. (2022). The non-essential TSC complex component TBC1D7 restricts tissue mTORC1 signaling and brain and neuron growth. Cell Rep 39, 110824. 10.1016/j.celrep.2022.110824.

60. Bishop, G.A., Berbari, N.F., Lewis, J., and Mykytyn, K. (2007). Type III adenylyl cyclase localizes to primary cilia throughout the adult mouse brain. J Comp Neurol 505, 562–571. 10.1002/cne.21510.

61. Su, K.H., and Dai, C. (2017). mTORC1 senses stresses: Coupling stress to proteostasis. Bioessays 39. 10.1002/bies.201600268.

62. Heberle, A.M., Prentzell, M.T., van Eunen, K., Bakker, B.M., Grellscheid, S.N., and Thedieck, K. (2015). Molecular mechanisms of mTOR regulation by stress. Mol Cell Oncol 2, e970489. 10.4161/23723548.2014.970489.

63. Ma, Y., Vassetzky, Y., and Dokudovskaya, S. (2018). mTORC1 pathway in DNA damage response. Biochim Biophys Acta Mol Cell Res 1865, 1293–1311. 10.1016/j.bbamcr.2018.06.011.

64. Tang, Z., Lin, M.G., Stowe, T.R., Chen, S., Zhu, M., Stearns, T., Franco, B., and Zhong, Q. (2013). Autophagy promotes primary ciliogenesis by removing OFD1 from centriolar satellites. Nature 502, 254–257. 10.1038/nature12606.

65. Pampliega, O., Orhon, I., Patel, B., Sridhar, S., Diaz-Carretero, A., Beau, I., Codogno, P., Satir, B.H., Satir, P., and Cuervo, A.M. (2013). Functional interaction between autophagy and ciliogenesis. Nature 502, 194–200. 10.1038/nature12639.

66. Maharjan, Y., Lee, J.N., Kwak, S., Lim, H., Dutta, R.K., Liu, Z.Q., So, H.S., and Park, R. (2018). Autophagy alteration prevents primary cilium disassembly in RPE1 cells. Biochem Biophys Res Commun 500, 242–248. 10.1016/j.bbrc.2018.04.051.

67. Huangfu, D., Liu, A., Rakeman, A.S., Murcia, N.S., Niswander, L., and Anderson, K.V. (2003). Hedgehog signalling in the mouse requires intraflagellar transport proteins. Nature 426, 83–87. 10.1038/nature02061.

68. Gerdes, J.M., Liu, Y., Zaghloul, N.A., Leitch, C.C., Lawson, S.S., Kato, M., Beachy, P.A., Beales, P.L., DeMartino, G.N., Fisher, S., et al. (2007). Disruption of the basal body compromises proteasomal function and perturbs intracellular Wnt response. Nat Genet 39, 1350–1360. 10.1038/ng.2007.12.

69. Corbit, K.C., Shyer, A.E., Dowdle, W.E., Gaulden, J., Singla, V., Chen, M.H., Chuang, P.T., and Reiter, J.F. (2008). Kif3a constrains beta-catenin-dependent Wnt signalling through dual ciliary and non-ciliary mechanisms. Nat Cell Biol 10, 70–76. 10.1038/ncb1670.

70. Corbit, K.C., Aanstad, P., Singla, V., Norman, A.R., Stainier, D.Y., and Reiter, J.F. (2005). Vertebrate Smoothened functions at the primary cilium. Nature 437, 1018–1021. 10.1038/nature04117.

71. Haycraft, C.J., Banizs, B., Aydin-Son, Y., Zhang, Q., Michaud, E.J., and Yoder, B.K. (2005). Gli2 and Gli3 localize to cilia and require the intraflagellar transport protein polaris for processing and function. PLoS Genet 1, e53. 10.1371/journal.pgen.0010053.

72. Huangfu, D., and Anderson, K.V. (2005). Cilia and Hedgehog responsiveness in the mouse. Proc Natl Acad Sci U S A 102, 11325–11330. 10.1073/pnas.0505328102.

73. Niehrs, C., Da Silva, F., and Seidl, C. (2025). Cilia as Wnt signaling organelles. Trends Cell Biol 35, 24–32. 10.1016/j.tcb.2024.04.001.

74. Hilgendorf, K.I., Myers, B.R., and Reiter, J.F. (2024). Emerging mechanistic understanding of cilia function in cellular signalling. Nat Rev Mol Cell Biol 25, 555–573. 10.1038/s41580-023-00698-5.

75. Parish, C.R. (1999). Fluorescent dyes for lymphocyte migration and proliferation studies. Immunol Cell Biol 77, 499–508. 10.1046/j.1440-1711.1999.00877.x.

76. Sanjana, N.E., Shalem, O., and Zhang, F. (2014). Improved vectors and genome-wide libraries for CRISPR screening. Nat Methods 11, 783–784. 10.1038/nmeth.3047.

77. Stringer, B.W., Day, B.W., D’Souza, R.C.J., Jamieson, P.R., Ensbey, K.S., Bruce, Z.C., Lim, Y.C., Goasdoue, K., Offenhauser, C., Akgul, S., et al. (2019). A reference collection of patient-derived cell line and xenograft models of proneural, classical and mesenchymal glioblastoma. Sci Rep 9, 4902. 10.1038/s41598-019-41277-z.

78. Kaizuka, T., Morishita, H., Hama, Y., Tsukamoto, S., Matsui, T., Toyota, Y., Kodama, A., Ishihara, T., Mizushima, T., and Mizushima, N. (2016). An Autophagic Flux Probe that Releases an Internal Control. Mol Cell 64, 835–849. 10.1016/j.molcel.2016.09.037.

79. Wiederschain, D., Wee, S., Chen, L., Loo, A., Yang, G., Huang, A., Chen, Y., Caponigro, G., Yao, Y.M., Lengauer, C., et al. (2009). Single-vector inducible lentiviral RNAi system for oncology target validation. Cell Cycle 8, 498–504. 10.4161/cc.8.3.7701.

