## Supplemental figures for "Growth Factor-Independent mTORC1 Signaling Promotes Primary Cilia Length via Suppression of Autophagy"

### SUPPLEMENTAL FIGURE LEGENDS

**Supplemental Figure 1. Primary cilia formation depends on the cell cycle, while mTORC1 signaling regulates primary cilia length dynamics independently of the cell cycle and protein abundance.**
(A) Flow cytometric analysis of cell cycle distribution via genomic incorporation of EdU as a function of DAPI-based nuclear intercalation in RPE1 cells over a time course of serum starvation.
(B–C) (B) Primary cilia length and (C) ciliation percentage of RPE1 cells over the course of serum starvation.
(D) Histogram of flow cytometric analysis showing cell population distribution based on G0 phase reporter levels (mVenus-p27$K^{-}$) in the G0 reporter RPE1 cells during serum starvation.
(E) Immunoblot of RPE1 cells treated with DMSO, FBS, or rapamycin.
(F) Primary cilia length of RPE1 cells treated with rapamycin in three different commercially available media, as indicated.
(G) Histogram of flow cytometric analysis showing cell population distribution by mVenus-p27$K^{-}$ levels in the G0 reporter RPE1 cells treated with DMSO, FBS (10%), rapamycin, or Torin1.
(H) Histogram of flow cytometric analysis showing cell population distribution based on protein abundance via CFSE labeling in RPE1 cells treated with DMSO, FBS (10%), rapamycin, Torin1, or cycloheximide. Live cells were incubated with CFSE (10 μM) for 1 h prior to flow cytometry.
All treatments were applied for 24 hours in serum-free conditions (48 h total):
DMSO (0.1%), rapamycin (20 nM), Torin1 (250 nM), cycloheximide (200 uM).
Statistical analysis was performed using two-way ANOVA followed by Šídák's multiple comparisons test. The significance threshold (α) was set at 0.05, and all p-values are reported.

**Supplemental Figure 2. Deletion of TSC complex components causes TP53-dependent cell cycle arrest, while TP53 deletion has no effect on primary cilia length dynamics.**

(A) Flow cytometric analysis of cell cycle distribution via genomic incorporation of EdU as a function of DAPI-based nuclear intercalation in RPE1 cells with deletion of endogenous TSC1 (sgTSC1) or TSC2 (sgTSC2), alone or in combination with TP53 deletion (sgTP53).

(B) Immunoblot of parental and sgTP53 RPE1 cell lines treated with actinomycin D (1 µg/mL) for 3 h.

(C) Immunoblot of parental and sgTP53 RPE1 cells subjected to serum or amino acid deprivation overnight, followed by re-addition for 30 minutes. Proteins of interest indicated by black arrowheads.

(D–E) (D) Primary cilia length and (E) ciliation percentage in parental and sgTP53 RPE1 cell lines. was used for comparisons between two groups.

Statistical analysis was performed using Unpaired two-tailed Student's t-test. The significance threshold (α) was set at 0.05, and all p-values are reported.

**Supplemental Figure 3. Protein expression in sgULK1 and sgBECN1 cell lines, and effects on autophagy in TSC-null cell lines.**

(A) Immunoblot of parental, sgULK1, and sgBECN1 RPE1 cell lines.

(B–C) (B) Flow cytometric analysis and (C) quantification of autophagic flux in parental, sgTSC1, sgTSC2, and sgTBC1D7 RPE1 cell lines treated with vehicle or Torin1 (250 nM) for the final 3 h of an overnight serum starvation using the reporter described in Figure 3A.

Statistical analysis was performed using two-way ANOVA followed by Šídák's multiple comparisons test. The significance threshold ($\alpha$) was set at 0.05, and all p-values are reported.

**Supplemental Figure 4. Generation of *Tbc1d7*$^{+/-}$ and *Tbc1d7*$^{-/-}$ embryos for neuron culturing.**
(A) Schematic diagram of the mouse *Tbc1d7* gene locus with the sgRNA target site indicated by an arrowhead.
(B) Breeding strategy schematic used to generate *Tbc1d7*$^{+/-}$ and *Tbc1d7*$^{-/-}$ littermate embryos.
(C) Representative DNA gel electrophoresis analysis showing *Tbc1d7* genotypes of embryos obtained from the breeding described in (B).

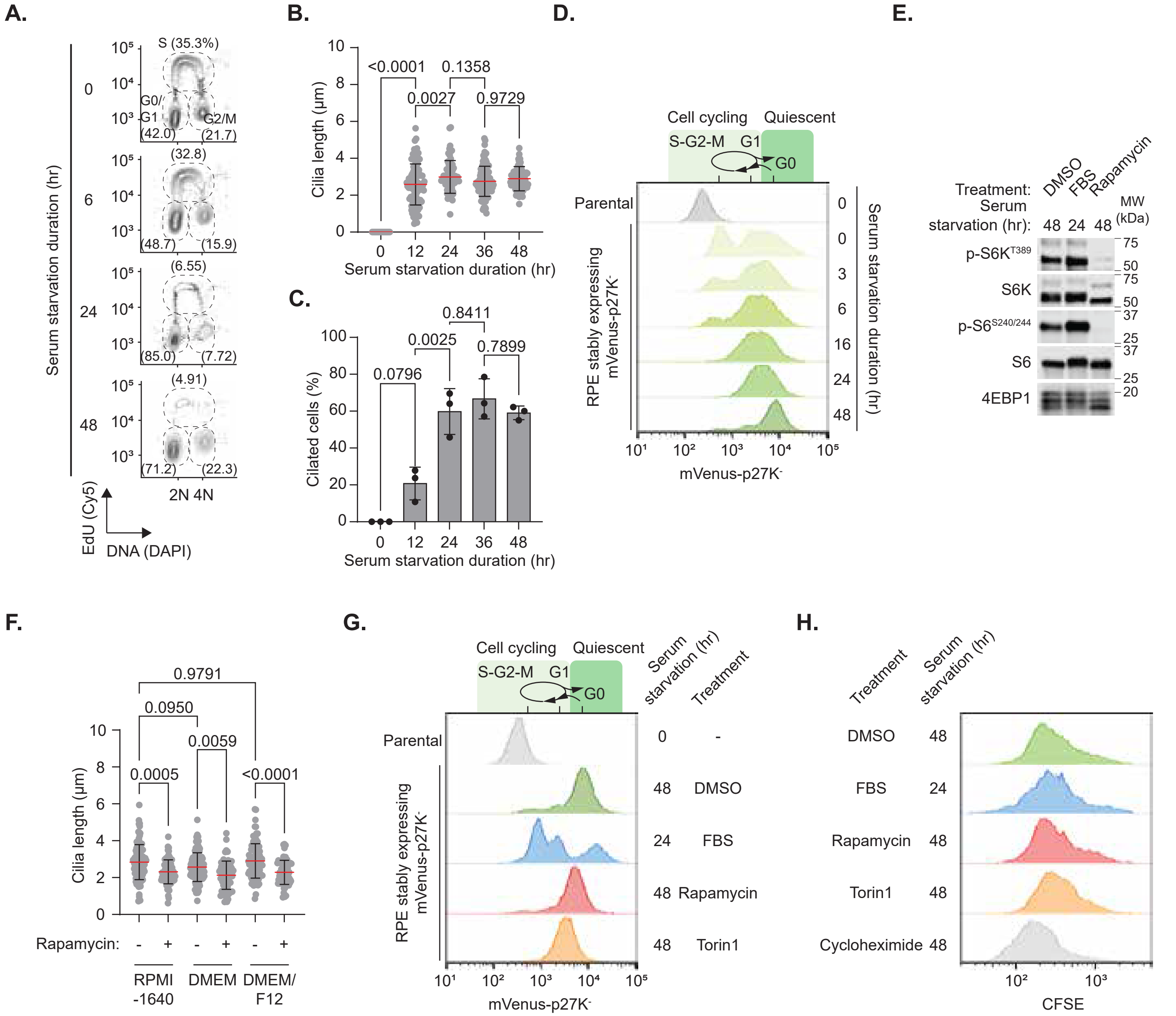


Supplemental Figure 1

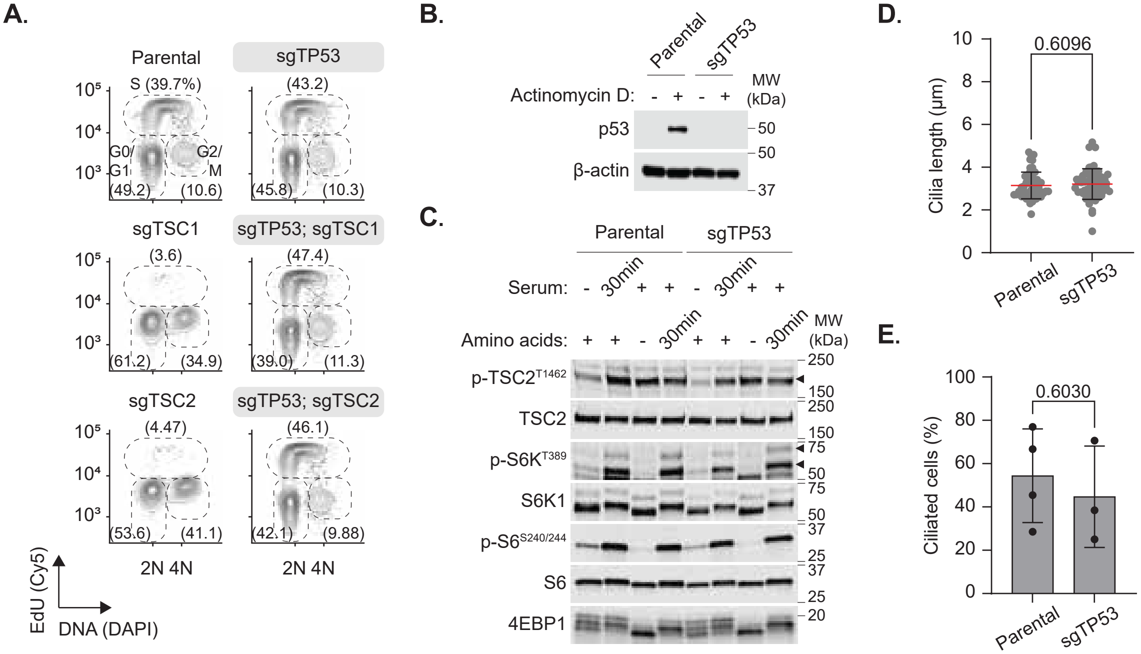


Supplemental Figure 2

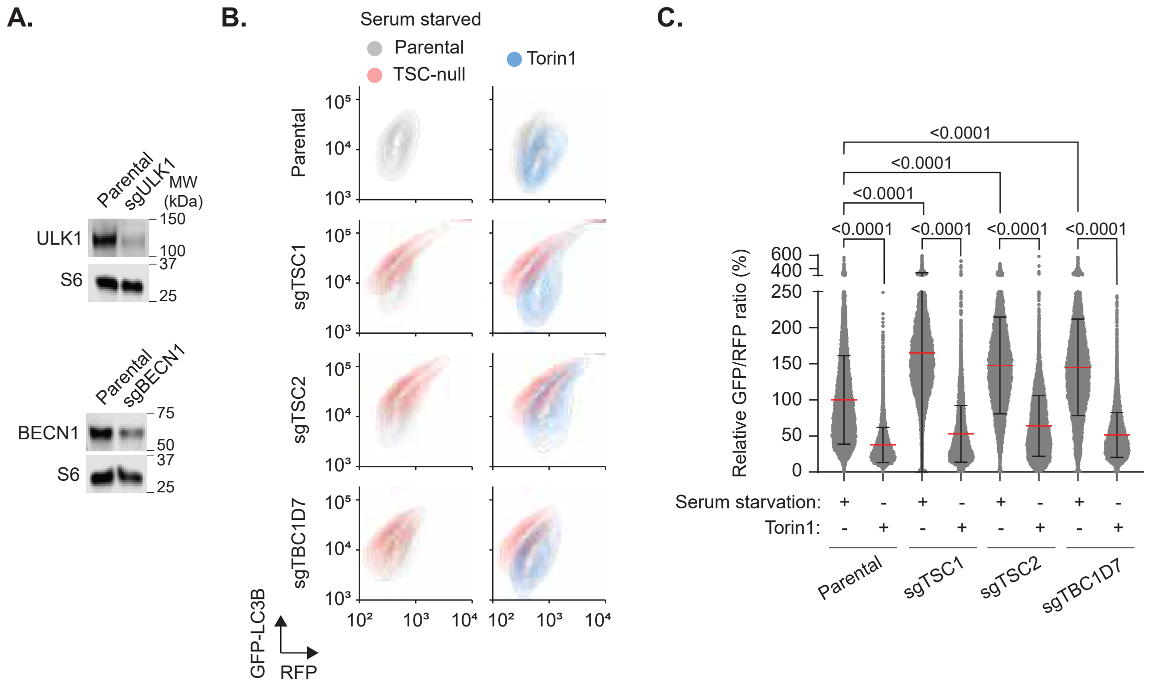


Supplemental Figure 3

**A.**

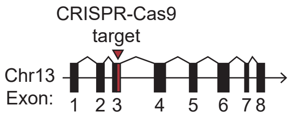


**B.**

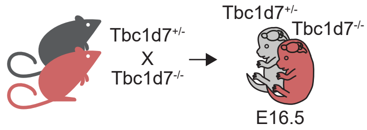


**C.**

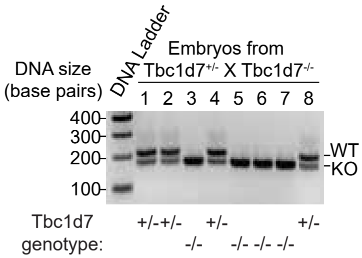


Supplemental Figure 4
